# The Fusiform Gyrus Processes Faces Relative to an Overall Face Average and a Person Average

**DOI:** 10.1101/861625

**Authors:** Zarrar Shehzad, Eunjoo Byeon, Gregory McCarthy

## Abstract

We are highly accurate at recognizing familiar faces even with large variation in visual presentation due to pose, lighting, hairstyle, etc. The neural basis of such within-person face variation has been largely unexplored. Building on prior behavioral work, we hypothesized that learning a person’s average face helps link the different instances of that person’s face into a coherent identity within face-selective regions within ventral occipitotemporal cortex (VOTC). To test this hypothesis, we measured brain activity using fMRI for eight well-known celebrities with 18 naturalistic photos per identity. Each photo was mapped into a face-space using a neural network where the Euclidean distance between photos corresponded with face similarity. We confirmed in a behavioral study that photos closer to a person’s average face in a face-space were judged to look more like that person. fMRI results revealed hemispheric differences in identity processing. The right fusiform face area (FFA) encoded face-likeness with brain signal increasing the closer a photo was to the average of all faces. This suggests that the right FFA pattern matches to an average face template. In contrast, the left FFA and left anterior fusiform gyrus (aFus) encoded person-likeness. The brain signal increased the further a photo was from the person’s average face weighted by the features most relevant for face identification. This suggests that the left FFA and aFUS processes an identity error signal. Our results encourage a new consideration of the left fusiform in face processing, specifically for within-person processing of face identity.

## Introduction

We are easily able to recognize familiar faces in a variety of contexts, even in very blurry or noisy photos (Jenkins & Burton, 2011). In contrast, we experience difficulties identifying faces of unfamiliar individuals (Johnston & Edmonds, 2009; Burton et al., 2011). When matching pairs of unfamiliar faces, accuracy and reaction-time is similar to matching pairs of inverted faces, whereas matching familiar faces show much higher accuracy and faster reaction-times (Megreya & Burton, 2006). Furthermore, different photos of the same unfamiliar person’s face are more likely to be interpreted as different people than photos of the same familiar face (Jenkins et al., 2011). These findings suggest a transformation from a noisy image-based processing for unfamiliar faces to selective person-based processing for familiar faces.

Previous studies explain familiar face processing as a process of comparing in reference to a face prototype or average face. An average of all faces is located in the center of the face-space, and each face is represented by its direction and distance to this center-average (Valentine, 1991; Valentine et al., 2016). Different directions from the average indicate different facial identities, and increasing geometric distance from the average corresponds to greater atypicality or distinctiveness of a face. Unfamiliar faces and those from other races are represented further away from the center-average (Valentine, 1991). The dimensions of face-space are not tuned to recognize features of those faces, making it challenging to discriminate among unfamiliar faces. As a consequence, unfamiliar facial identities will clump together in face-space. Learning a face is then the process of discerning the distinctive features of an identity in reference to the center-average of all faces.

The center-average model ignores the within-identity variation of the visual presentations of the same person, such as pose, lighting, expression, and age. Each point in face-space is considered to be a distinct identity and any alteration in face-space results in a change in identity. Such challenge is evident in studies of unfamiliar faces. For example, variance in social impressions of the same unfamiliar individual has been found to be as large as variation between individuals (Jenkins et al., 2011; Todorov & Porter, 2014; Sutherland et al., 2016). Furthermore, Jenkins and colleagues (2011) found that photos of the same unfamiliar person were often recognized as different identities, while photos of different people were never recognized as the same identity. These findings show that between-identity recognition was good, but within-identity recognition was poor for unfamiliar faces.

Moreover, when participants were familiar with the face, there was significant improvement in within-identity face recognition, but not between-identity. However, the mechanism for how within-identity recognition improved was left unanswered.

A person-specific prototype model may provide an answer by linking the disparate visual representations of the same person together into one average face of that person. This prototype can be implicitly learned and represented as an average of all previous experiences with that face (Burton et al., 2005; 2011; Jenkins & Burton, 2011). This averaging process extracts regularity from multiple face images by removing information irrelevant to face identity (e.g., pose or lighting). Behavioral work supports the importance of such person-specific average. After seeing two exemplars of the same person, participants falsely remembered seeing an average representation of the two exemplars above chance (Kramer et al., 2015). In addition, the average face of a person was judged to look more like that person than any exemplar (Bruce et al., 2002; Frowd et al., 2014), and matching an average face to the person’s name was faster and more accurate than an exemplar (Burton et al., 2005). These findings suggest that we may store face averages of each familiar identity to process within-person variations, in addition to the center average of all faces.

A specialized brain network dedicated to the perceptual processing of faces encodes face prototypes (Gobbini & Haxby, 2007). Within this face network, brain activity for static faces (e.g., photos) has been localized to the ventral visual cortex and lateralized to the right hemisphere (Allison, Ginter et al., 1994; Kanwisher et al., 1997; McCarthy et al., 1997; Engell & McCarthy, 2013). Greater brain activity has been found for familiar than novel faces in the fusiform face area (FFA), the anterior fusiform (aFus), and the ventral anterior temporal lobe (vATL) (Von Der Heide et al., 2013). In addition, studies have been able to decode face identity using patterns of brain activity in the right vATL and, in some studies, the FFA (Kriegeskorte et al., 2007; Nestor et al., 2011; Verosky et al., 2013; Anzellotti et al., 2014; Axelrod & Yovel, 2015). On the other hand, the neural response for unfamiliar faces codes each face identity by its deviation from the center average (Loffler et al. 2005; Mattar et al. 2016). We may expect that familiar faces should also be represented relative to the center average in face-selective regions of the right hemisphere.

Previous fMRI studies have addressed the neural representation of between-person identity, seeing the faces of different people, but not the neural representation of within-person identity, seeing different faces of the same person. Given prior behavioral work, we hypothesized that the person average, and not the center-average, will reflect the neural representation of within-person identity. The abstraction of a within-person identity into an average is likely associated with prototype-based representations in the left hemisphere. A study demonstrates that individuals can classify previously unseen prototypes of newly learned objects more efficiently when they are presented to the left hemisphere (right visual field) than to the right hemisphere (Marsolek, 1995). In addition, within-person identity is associated with other processes, such as person knowledge and language, that are localized to the left hemisphere including the FFA and vATL (Gainotti, 2013; Rice et al., 2015). We may expect activity in the face-selective regions of the left hemisphere for processing familiar faces relative to a person average, in addition to the activity in the right hemisphere for center average.

Here, we test if neural activity associated with face processing reflects between-person identity, represented by deviations from the center average, and within-person identity, represented by deviations from the person average. We presented photos of eight celebrities to participants while fMRI scans were acquired. Participants were required to press a button corresponding to the celebrity shown on the screen. The photos were represented in a face-space using a neural network previously trained on 500,000 images to compare identities of different faces (Schroff et al., 2015; Amos et al., 2016). Each photo in our experiment was mapped onto a 128-dimensional Euclidean space using this trained network. We used independent datasets of photos containing 1,500 images to generate the center average across all faces and the person average for each celebrity. Then we measured the similarity of each photo to the center average and to the person average. We used these measures to assess if the similarity to the center or person average was behaviorally meaningful and corresponded to either response times or likeness ratings (i.e., how much a photo looks like the person). Then, we assessed if the similarity of each photo to the center or person average is related to brain activity in regions of interest (ROIs) along the ventral visual cortex, including the FFA and the vATL. Since prior work has suggested that not all face features are equally relevant in person recognition (Ellis et al., 1979; Young et al., 1985; Clutterbuck & Johnston, 2002), we investigated if behavioral and neural models are improved with distances weighted by features that are the most important in identifying a person’s face. We predicted that distances to the center average would be localized to the right fusiform, supporting specialization of the right hemisphere for face perception. In contrast, distances to the person average would be localized to the left fusiform, supporting specialization of the left hemisphere for person recognition and naming.

## Methods

### Participants

Six right-handed, healthy adults (5 females) participated in the study and each participated in four experimental sessions. All had normal or corrected-to-normal vision and no history of neurological or psychiatric illnesses. The protocol was approved by the Yale Human Investigation Committee, and informed consent was obtained for all participants.

### Stimuli

We identified 24 celebrities (12 males; 12 females) that were well-known to all participants. We collected photos of each celebrity from the FaceScrub (Ng & Winkler, 2014) and PubFig (Kumar et al., 2009) datasets supplemented by Google Image search. We accepted photos in which a face could be detected and landmarks (e.g., nose, eyes) could be accurately identified using an automated algorithm in the dlib toolbox (King, 2009). The selected photos for each identity had considerable image variability reflecting natural variation in pose, luminance, expression, content, age, etc. Figure 1 shows the variability in the photos of one celebrity.

**Figure 1:**
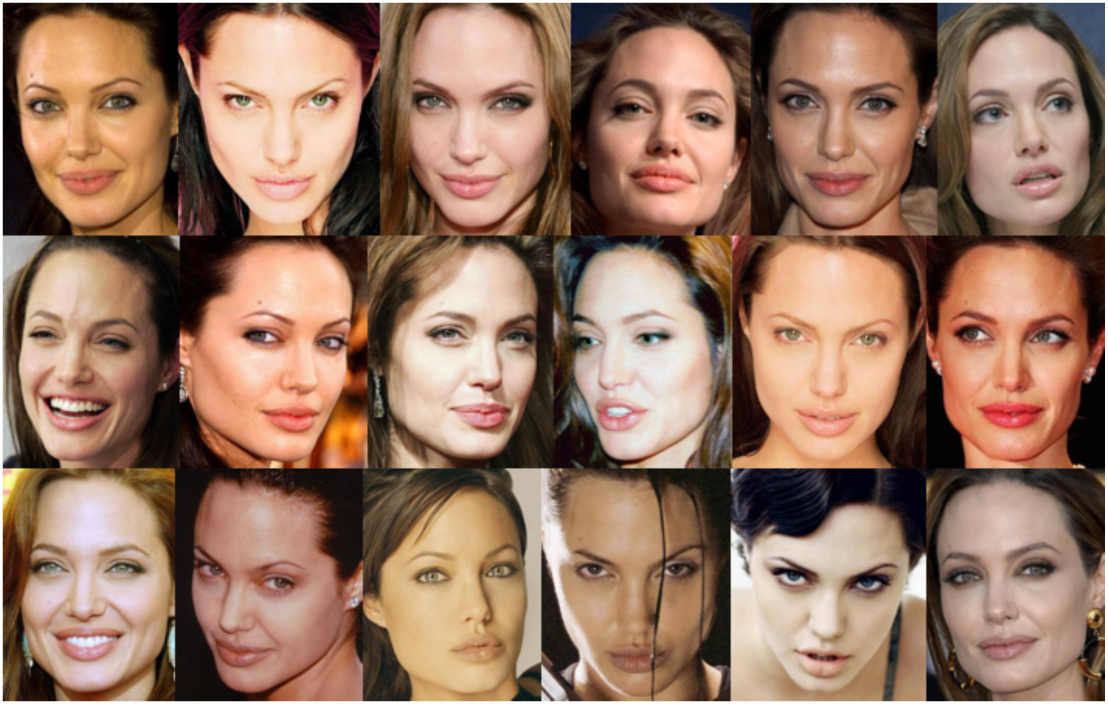
18 photos that were used in the experiment for one of the celebrities, Angelina Jolie.

We selected a subset of photos to be used in the experiment and other photos to be used in subsequent analyses for calculating person-specific face averages. In this experiment, eight celebrities served as targets and 16 celebrities served as distractors. Eighteen photos were shown for each target stimulus, while two photos were shown for each distractor stimulus. Supplementary Table 1 provides the names of each celebrity. In later analyses, we used an additional 120-180 photos for each target celebrity. Distractor stimuli were not used in later analyses.

Additionally, we collected an independent dataset of 864 short videos from YouTube. Each video was of a different person unfamiliar to participants. Eight evenly spaced frames showing different views of the same person from each video were selected and used to calculate the center average.

### Procedure

Each participant was studied in four visits. In each visit, participants completed several tasks during a functional magnetic resonance imaging (fMRI) scan. Here, we only report on one task that participants completed and the other two tasks involving dynamic face videos will be reported later.

Participants viewed faces and scenes in a rapid event-related design (Figure 2). Each trial began with a photo of a face or scene displayed on the screen for 1s followed by a 3s fixation. Photos of scenes compromised 25% of trials. On a given face trial, participants saw one of the target celebrities (18 exemplars each), or occasionally on 10% of face trials, one of the distractor celebrities (2 exemplars each). Participants saw half of the celebrities (4 targets and 8 distractors) in the first two scan visits and the other half of celebrities in the last two visits. During a scan, they were instructed to press a button corresponding to each of the four target celebrities displayed on the screen using both hands (two celebrities were associated with each hand), and to withhold a response for a distractor face. Each trial (face and scene) was repeated four times.

**Figure 2:**
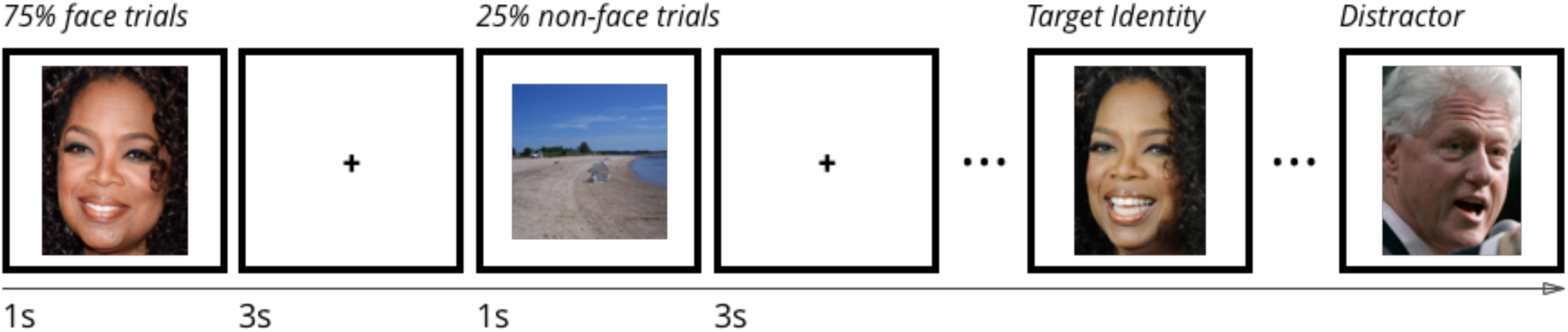
Participants saw photos of 8 celebrities (e.g., Oprah Winfrey) with 18 exemplars for each celebrity. Each photo was repeated four times. On a given run, participants saw 4 of the 8 celebrities and pressed a button corresponding to the person shown. Participants did not press a button when scenes were shown or when one of the 16 distractor faces were shown.

### Likeness Ratings

Given the variability in our images, some photos capture a person’s identity better than others. To quantify this variability in person identity, we asked participants to rate how much each photo looked like that person. Following each fMRI scan, participants completed an online rating of photos for the eight target celebrities. For each photo, participants were asked to provide a likeness rating using a 7-point Likert scale, where 1 indicated an extremely poor likeness, and 7 indicated an extremely good likeness. There was no time limit for the task, and each image stayed on the screen until a response was made.

### Computational Measures

#### Face representations in a neural network

We transformed each photo into an abstract face representation using a deep neural network. All steps were carried out in Python. First, we detected the location of the face using dlib as part of the menpo package (King, 2009; Alabort-i-Medina et al., 2014). Then we used annotations of the eyes and nose to align each face to an average template. Finally, we input the aligned image into the OpenFace neural network, which outputs a 128-dimensional face representation (Amos et al., 2016). Figure 3 shows a 2-dimensional projection of the face representations of all target stimuli using the *t*-sne algorithm.

**Figure 3:**
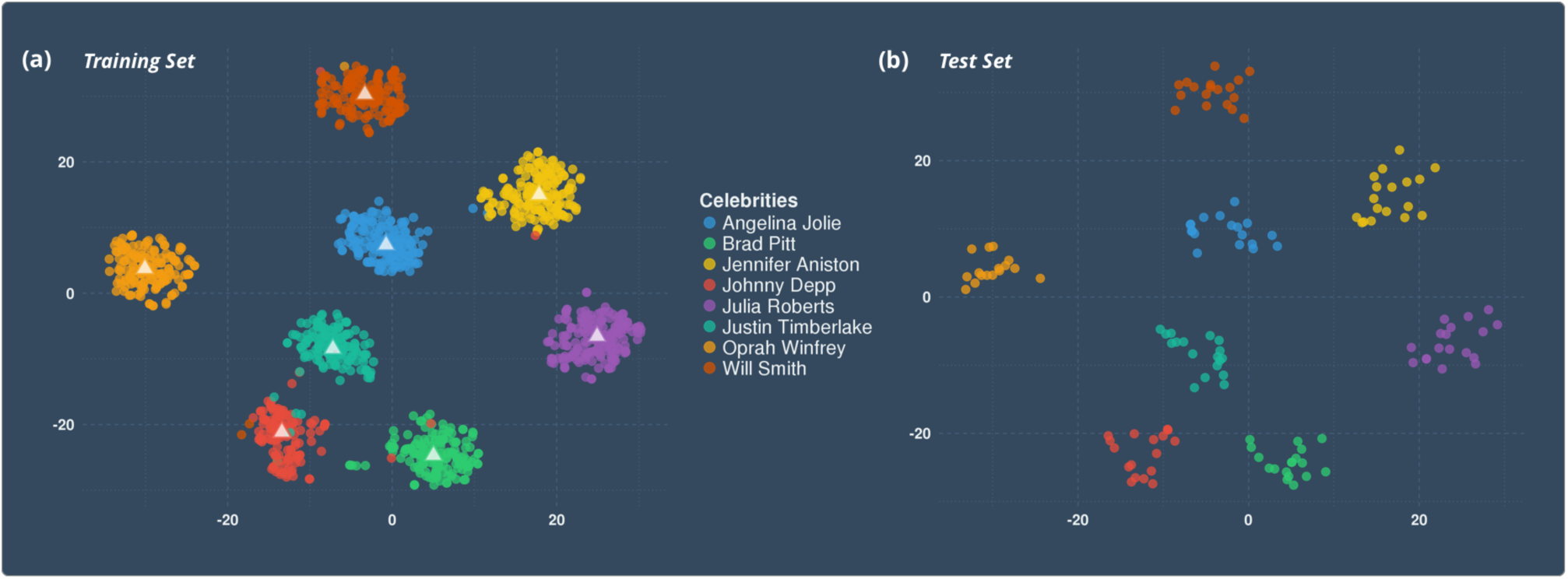
Representations of each photo (shown as circles) in the 128-dimensional neural network space projected onto 2-dimensions using t-sne. Each color indicates a different celebrity. (a) On the left are photos from the training set, which were used to generate the average faces for each person (shown as triangles). (b) On the right are photos from the test set, which were shown to participants in the experiment.

The output of OpenFace reflects a common notion of face-space. OpenFace was trained with 500,000 images to determine if a pair of images were from the same person or from different people. Consequently, each point represents a face in Euclidean space and distances between points directly correspond to a measure of face similarity. Faces closer together tend to look more similar and be of the same identity. We found that OpenFace’s output represented face features such as age and attractiveness, while being pose and illumination invariant (Supplementary Materials; Supplementary Figure 1). Since these qualities are similar to behavioral descriptions of face-space (Valentine, 1991; 2001; Valentine et al., 2016), we concluded that the outputs of OpenFace were appropriate for our analyses.

#### Face prototypes

We measured each photo in relation to the person average or the center average (Figure 4). All averages were taken after transforming the image data to the 128-dimensional face-space. First, to generate a face prototype, we took the average of all faces in the 864 video dataset (8 frames per video). This overall average tends to be in the center of face-space and also referred to as the center average. We calculated the Euclidean distance of each photo of celebrities to this center average. Photos closer to the center average were considered as being more face-like. Second, to generate a person prototype, we used photos from the training set and took the average of 120-180 faces for each target celebrity. We calculated the distance of each photo to that person’s average face. Photos closer to a person’s average were considered to look more like that person. By taking the center and person averages from larger datasets independent of the faces shown in the experiment, we hoped to better represent the veridical face averages.

**Figure 4:**
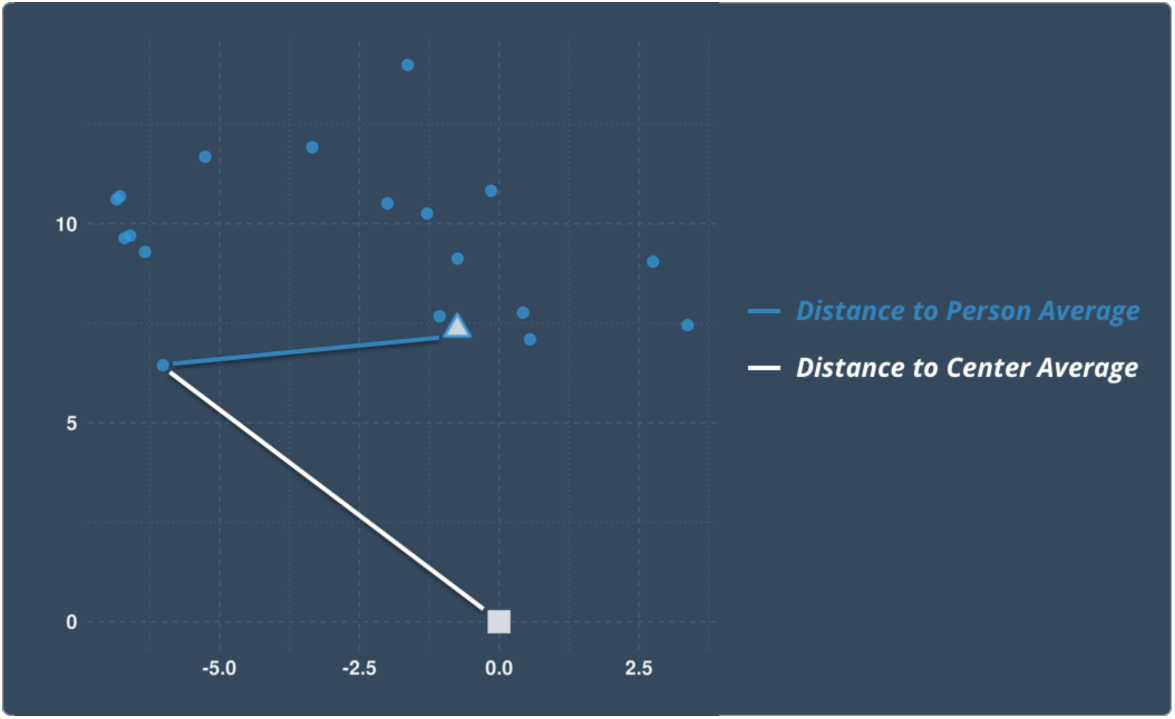
Photos for Angelina Jolie (circles) are shown in two-dimensions along with her average face (triangle) and the average of all faces in the center of face space (white square). Two face space measures are depicted: the distance of a photo to the center average (white line) and the distance of a photo to the person average (blue line).

### Behavioral Analysis

All behavioral analyses on likeness ratings and response times were conducted in the R statistical package (version 3.2.3). We conducted two-way analysis of variance (ANOVA) on likeness ratings and response times. We included the person-identity as a factor along with the distance to the center and person average as continuous measures.

### Image Acquisition and Preprocessing

Data were acquired using a 3T Siemens Prisma scanner with a 64-channel head coil. Functional images were acquired using a multiband echo-planar pulse sequence (TR = 1000 ms, TE = 27.8 ms, flip angle = 60°, multi-band acceleration factor = 5, GRAPPA acceleration factor = 2, FOV = 220 × 220 mm, matrix = 110 × 110, slice thickness = 2 mm, 75 slices, voxel size = 2.0 mm^3^). A high-resolution structural image was acquired for registration using a 3D MP-RAGE sequence (TR = 1900 ms, TE = 2.52 ms, flip angle = 9°, FOV = 256 × 256 mm, matrix = 256 × 256, slice thickness = 1 mm, 208 slices).

Image preprocessing was performed using custom scripts available at https://github.com/HumanNeuroscienceLab/face_representations. Further details about the preprocessing pipeline are provided in Supplementary Materials.

### Regions of Interest (ROI)

We generated regions of interest to examine the brain response associated with our average face measures in face-selective regions. We identified regions in the occipital face area (OFA), the fusiform face area (FFA), the anterior fusiform (aFus) and the ventral anterior temporal lobe (vATL) based on the Atlas of Social Agent Perception (Engell & McCarthy, 2013; Supplementary Materials). We also included a control ROI (primary visual cortex, V1) in which we did not expect any face-related effects.

### fMRI Data Analysis

We conducted selected region-of-interest and whole-brain voxelwise analyses (Supplementary Materials). In our principal analyses, we examined associations between brain activity and the distance of each photo to the overall average face and the person average face. To see if facial features are scaled by how well they discriminate between identities, we considered both unweighted and weighted distances to the person average. Across all ROIs except V1, the weighted distances showed a stronger association with brain activity (Supplementary Materials). Here, we report the analyses using the weighted distances to the person-average. Finally, we examined if the association between distance to each average face and brain activity is behaviorally meaningful and mediated by likeness ratings or response times. For each analysis, we included several covariates of non-interest: baseline effects of each run, six head movement parameters, incorrect trials, no response trials, and amplitude-modulated response time to account for the effect of button presses.

Additional analyses sought to characterize the sensitivity of regions to familiar faces and to the behavioral measures. We identified regions selective for faces overall (contrast of face > scene activity) and regions selective for face familiarity (association with the average response time for each identity). Finally, we measured the association between brain activity and behavioral measures of person identity (likeness ratings) and face recognition (response times).

## Results

### Behavioral Results

Response time (RT) significantly varied by the celebrity shown (main effect of identity; F_7,134_ = 21.8, *p* < 0.05), while the likeness ratings only marginally differed by the celebrity shown (F_7,134_ = 2.0, *p* = 0.06). When examining how well the two face-space measures (distance of each photo to the center average, and to the appropriate person average) explained RTs and likeness ratings, we found a significant effect of distance to the person average on likeness ratings (t_134_ = -4.6, *p <* 0.05), but no effect was found for any other comparison (all *ps* > 0.05; Figure 5). Regressors for person identity as well distances to the center average and the person average were included together in all models tested. Supplementary Table 1 presents descriptive statistics of each celebrity for the two participant measures (likeness ratings and RTs) and for the two, quantitative face-space measures.

**Figure 5:**
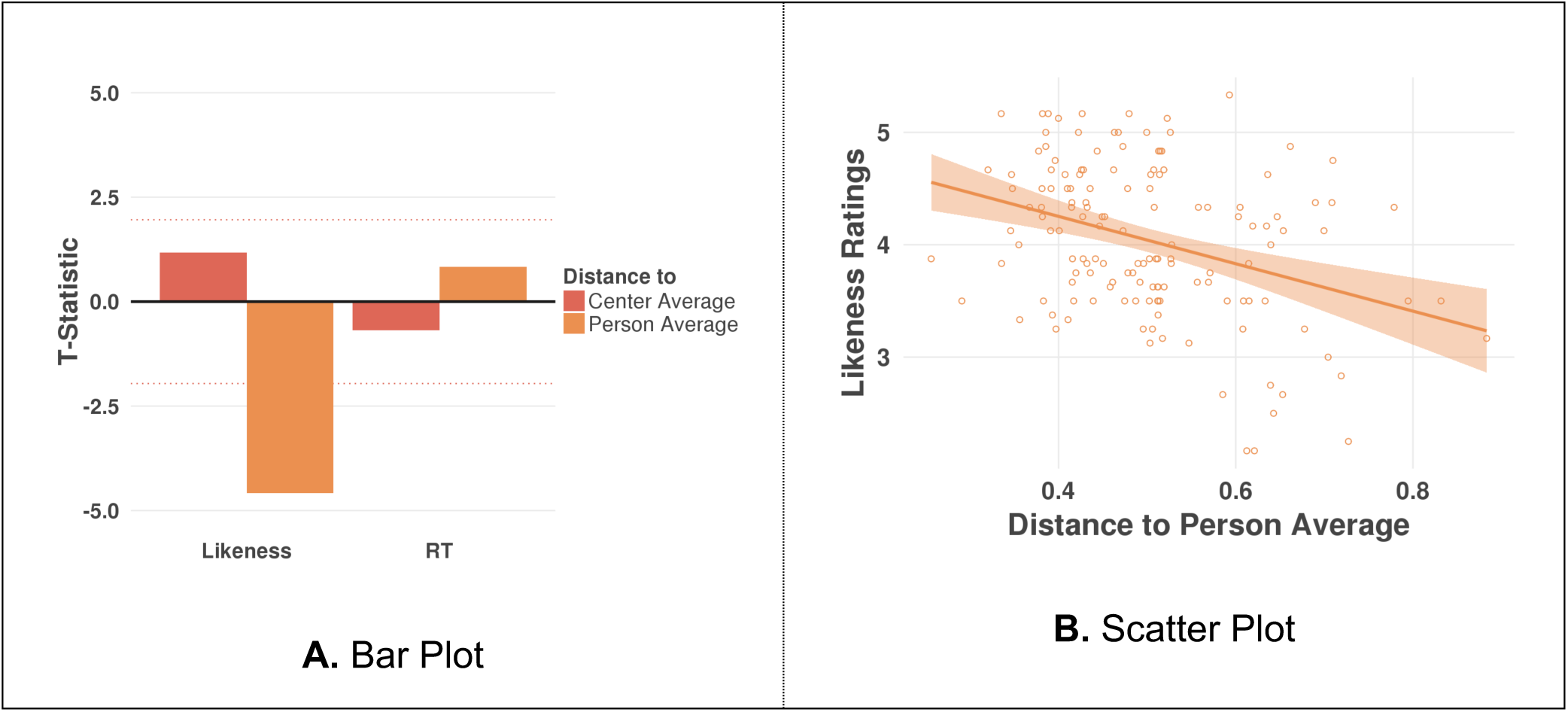
(a) Distances between each photo and the center average (red) or person average (orange) were used to explain variation in likeness ratings and response times (RT) along the x-axis. The t-statistic of the model fit is given on the y-axis. The dotted red lines indicate significance estimates (*p* < 0.05, two-tailed). (b) Scatter plot shows each photo’s (circle) Euclidean distance to its person average (x-axis) and the group-average likeness rating of that photo (y-axis). The regression fit is shown as the solid line with standard errors depicted as the surrounding shaded ribbons. The main effect of person identity and distance to the center average were regressed out prior to making this plot.

**Figure 6:**
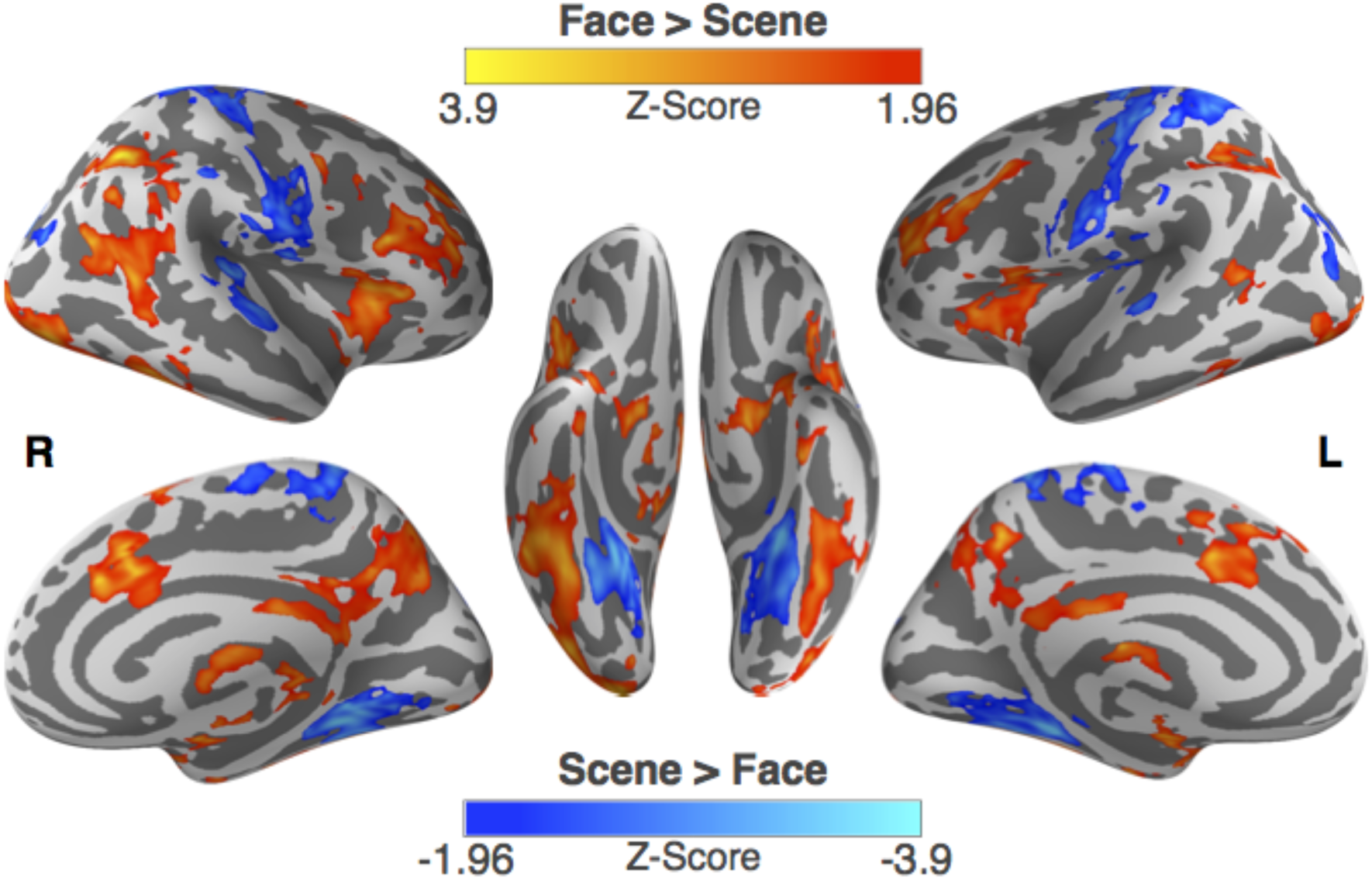
Brain activity for photos of faces and scenes (*p* < 0.05, cluster-corrected). Significantly greater activity for faces versus scenes (red-yellow) and scenes versus faces (blue-cyan) is shown on the fsaverage surface.

### Face Activity

We sought to first identify regions that were face-selective in our task. This served as a manipulation check to see if our findings were consistent with prior work. We included all face trials with both target and distractor celebrities, which allowed us to disentangle the general face response to all celebrities from the response to target celebrities. In addition, the amplitude-modulated RT was included as a covariate to remove the effect of the button press. As expected for face greater than scene trials, we found significant activation in known face-selective regions such as the FFA, the OFA, the pSTS, and the inferior frontal gyrus (*p* < 0.05, cluster corrected; Red-Yellow; Supplementary Table 3a). Similarly, for scene greater than face trials, we found significant activation in known scene-selective regions such as the parahippocampal gyrus (*p* < 0.05, cluster corrected; Blue-Cyan; Supplementary Table 3b). In addition, we observed activity in motor-related regions such as the pre-central gyrus, which was inversely related to participants seeing a face. The spread of activity for scene > face trials was likely smaller than typically observed because scene trials occurred only 25% of the time.

### Distances to the Center Average and the Person Average

To understand how face-selective regions represent familiar faces, we calculated two prototype measures using our 128-dimensional face-space. We identified regions across the whole-brain that represent a face relative to the overall average face and to the average face of each celebrity. We only report those regions that also show activity for faces greater than scenes here (*Z* > 1.65, uncorrected) but list all significant peak activations in Supplementary Table 4 and Supplementary Table 5.

First, we measured overall face processing by taking the distance of each photo to the average of all faces (center average; Figure 7a). For distances to the center average, we found only negative associations with brain activity distributed across three clusters in the right hemisphere. In all clusters, brain signal increased when photos were more similar to the center average. Significant clusters included the FFA and the middle temporal gyrus as well as the lateral prefrontal cortex (rostral middle frontal) and the inferior parietal sulcus.

**Figure 7:**
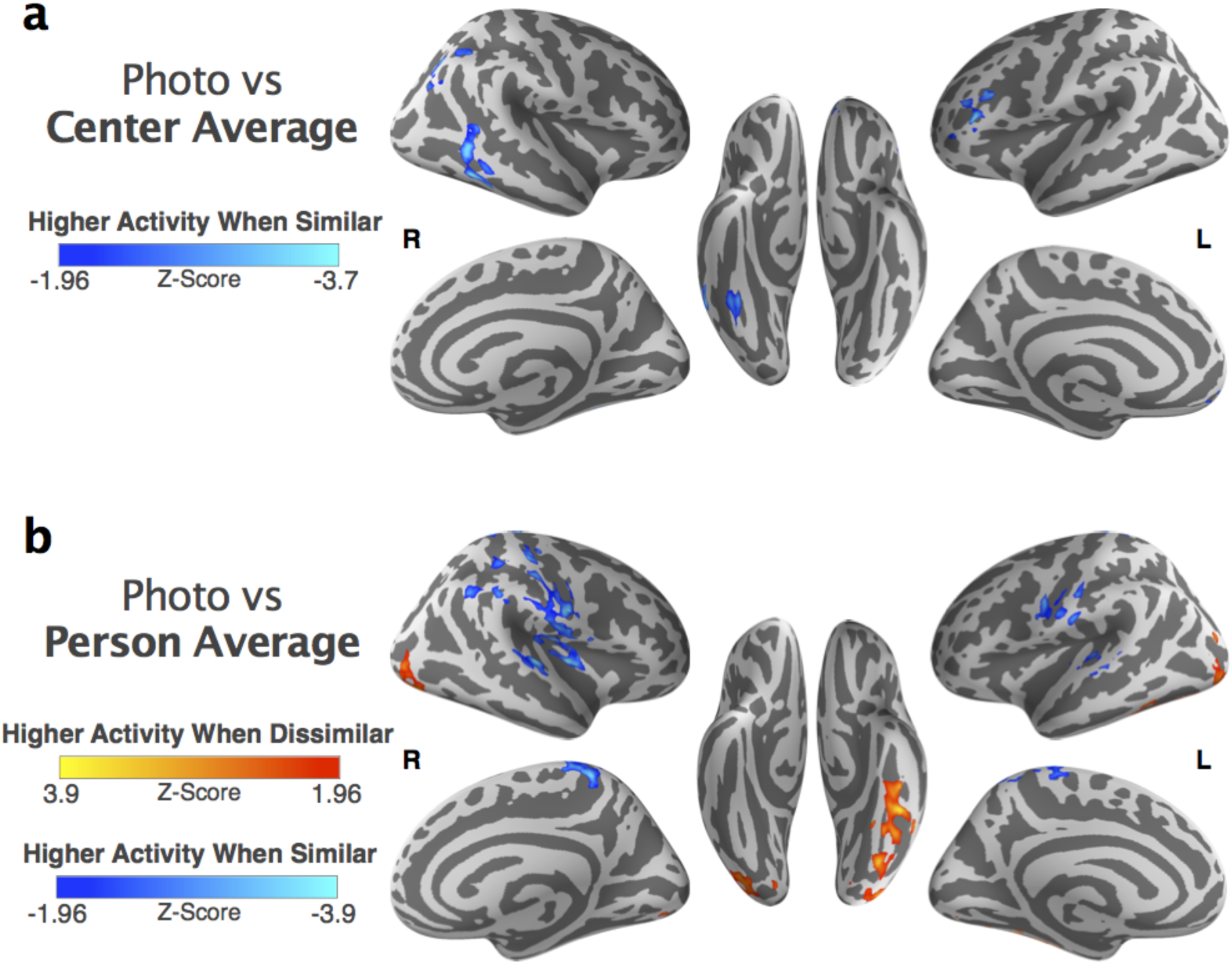
Brain activity is compared to the distance between each photo and average faces. Significant associations are displayed on the fsaverage surface (*p* < 0.05, cluster corrected). (a) Regions are shown where higher brain activity is associated with photos that are more similar to the center average (blue-cyan). (b) Below regions are shown where higher brain activity is associated with photos that are more dissimilar to the person average (red-yellow) or with photos that are more similar to the person average (blue-cyan). The distance to the person average was weighted by the most discriminable features using a multinomial ridge regression.

Second, we measured person-specific face processing by taking the <u>weighted</u> distance of each photo to the average of that person’s face (person average; Figure b). For distances to the person average, we found positive associations with brain activity in two clusters (i.e., brain signal increased when photos were less similar to the person’s average face). We found left-lateralized activity associated with the person average along the ventral visual cortex, stretching from the OFA to the FFA to the aFus. In contrast, the equivalent right hemisphere cluster in the ventral visual cortex only included the OFA and not more anterior regions. We also found negative associations of brain activity with the distances to the person average in three clusters (i.e., brain signal increased when photos were more similar to a person’s average face). However, the majority of these regions with negative association were not face-selective. The two face-selective regions negatively related to the person average were the inferior frontal gyrus and the supramarginal sulcus. These areas of the face network respond in a more categorical manner to a person (i.e., more activity if more like a person’s face), whereas in the fusiform, the inverse was true, more activity occurred the less a photo looked like a person’s face.

### Brain-Behavior Associations for the Person Identity

Finally, we sought to identify regions sensitive to subjective person identity, measured with likeness ratings (how much a face looks like a person) outside of a scanner. We identified regions across the whole-brain whose activity was associated with the likeness ratings. We only report regions that also show activity for faces greater than scenes (Z > 1.65, cluster-corrected) and list all significant peak activations in Supplementary Table 6 and 7. We found positive associations between brain activity and the likeness ratings distributed across five clusters (i.e., brain signal increased for faces that looked subjectively more like a person). However, all these regions with a positive association to likeness ratings were not face-selective except for a peak activation in the cerebellum (Figure 8, Red-Yellow). Instead, the negative associations of brain activity with likeness ratings were mostly found in face-selection regions (i.e., brain signal increased for faces that subjectively looked unlike the person) (Figure 8, Blue-Cyan). The results were primarily in the left anterior fusiform, a region that overlaps with the effects for the person average (Figure 7).

**Figure 8:**
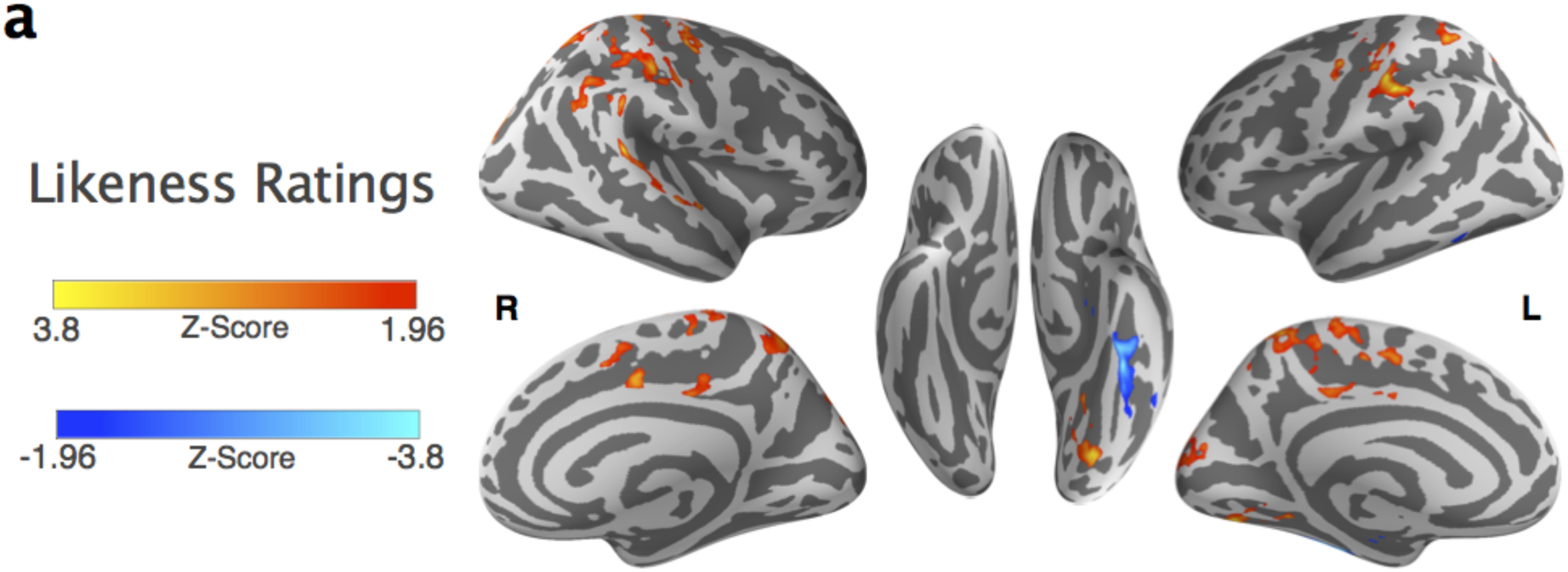
Significant brain-behavior associations are displayed on the fsaverage surface (*p* < 0.05, cluster corrected). Positive (red-yellow) and negative (blue-cyan) associations of brain activity with group-average likeness ratings are shown.

## Discussion

We found that famous faces are represented by two prototypes in face-space: the average of all faces (center average) and the average of a person’s face across many instances (person average). These prototype representations were localized to face-selective regions (OFA, FFA, and aFus) and showed pronounced laterality effects. The right hemisphere was more sensitive to a face prototype, responding more strongly to faces than non-faces, and responding more strongly to faces that looked more similar to the center average. In contrast, the left hemisphere was more sensitive to person-specific prototypes, showing greater activity for faces that looked less like the person average. Behaviorally, we found that the distance to the person average also reflected subjective likeness ratings: photos that reported to look more like a person were also closer in face-space to that person’s average face. Furthermore, the subjective ratings of likeness mediated the effect of distances to the person average on brain activity in face-selective regions (OFA, FFA, and aFus; see Supplementary Materials). Our findings suggest that seeing famous faces requires learning an average face, and such familiar faces are then represented within face-space in reference to both a face prototype (center average) and a person-specific prototype (person average).

### Center average

The right hemisphere is known to be specialized for face processing (Gazzaniga & Smylie, 1983; De Renzi et al., 1994; Le Grand et al., 2003; Yovel et al., 2008; Thomas et al., 2009). Our findings suggest that one element of face processing in the right hemisphere is matching a stimulus to a face template: the overall average face. Prior work has used face silhouettes that were manipulated for perceived face-likeness and found that the response in face-selective regions gradually increased with face-likeness (Davidenko et al. 2012), which is consistent with our findings. Similarly, the activity in the lateral occipital cortex increased when objects were more like the prototype (Iordan et al., 2016). In this notion, the distance to the center average may reflect a measure of face categorization and offer a continuous measure to the typically discrete contrasts found in localizers (e.g., face > scene). Thus, the right FFA engages in a process of pattern matching, constantly looking for faces in the world by matching inputs to an average face.

### Person average

While the distance to the center average may reflect a right lateralization of face recognition, the distance to the person average may instead reflect a left lateralization of within-person identity. Across behavioral, neurophysiological, and neuroimaging studies, the left hemisphere tends to be involved in the naming and categorization of familiar faces (Levy et al., 1972; Sergent, 1985; Ramon & Rossion, 2012; Gainotti, 2013; Rice et al., 2015).

Reflecting the importance of verbal processing in the left hemisphere, the left FFA is localized closely to the visual word form area sensitive to word processing (Cohen et al., 2000; Candliss, Cohen, & Dehaene, 2003; Puce, Allison, Asgari, Gore, & McCarthy, 1996; Nobre, Allison, & McCarthy, 1994). Previous studies have associated the left hemisphere with a prototype-based representation, potentially indicative of abstraction in word processing, and the right hemisphere with an exemplar-based representation (Marsolek, 1995). In addition, a reported preference for differing spatial frequencies in each hemisphere may be related to our findings (Sergent, 1985; Sergent & Holzer, 1982; Piazza & silver, 2017). Low spatial frequencies are processed relatively more efficiently in the right hemisphere, sufficient for face identification (i.e., is there a face or not), whereas high spatial frequencies are reported to be processed more efficiently in the left hemisphere facilitating person identification (i.e., who is this person). Since participants in the present study had to identify each celebrity with a button-press, they may have relied on the left hemisphere and associated verbal strategies for person identification. Participants may have used the name of each person as a cue to retrieve a person prototype, the average face of that person, that they then used to match to each photo.

Our work suggests that learning the average face of a person helps constitute the within-identity representation. In particular, our findings bring together two strands of behavioral work. On one hand, the average face has been associated with a person’s identity by providing an abstract and stable representation of the person (Burton et al., 2005; Jenkins & Burton, 2011). On the other hand, the variability among photos of the same individual has become better understood. Social judgements and identification performance for photos has been observed to be as large within-person as between-person (Jenkins et al., 2011; Todorov & Porter, 2014). Here, we found that both subjective and neural ratings of how much a photo looked like a person was explained by the distance between the photo and the person’s average face. Photos that were more similar to a person’s average face (lower variation) led to decreases in brain activity, reflecting more elaborative processing that is associated with distinctive faces. Taken together, our findings show the average face of a person acts as a reference point and links exemplars together to form a person representation localized in the left hemisphere.

### Weighted person average

In additional analyses (see Supplementary Materials), we found that the person prototype is not a simple average face as previously proposed (Burton et al., 2005; Jenkins & Burton, 2011), instead it is an average face that is scaled by the most discriminable features of that identity. Prior behavioral work has proposed that unfamiliar faces such as those of other-ethnicities tend to clump together in face-space due to suboptimal tuning of the dimensions in face-space, which then makes discrimination between unfamiliar faces difficult (Valentine & Endo, 1992; Chiroro & Valentine, 1995). Perceptual learning associated with a new person or a set of faces will then lead to optimization of the dimensions in face-space to more easily discriminate between the faces experienced. Our work supports such perceptual learning for familiar faces by demonstrating that each famous person had specific face features tuned to that identity and using the tuned face features significantly improved model fits across the ventral visual cortex.

Such tuning may reflect learning of internal face features (e.g., eyes, nose, mouth), which are important in developing an invariant face representation. As a face becomes more familiar, the internal features, and not the external features (e.g., hair), become more important in face identification (Ellis et al., 1979; Young et al., 1985; Clutterbuck & Johnston, 2002). In addition, the learned internal features tend to be processed in the left hemisphere (Haan & Kollenburg, 2005). Internal features like noses tend to be fairly stable across time while external features like hair can change from day-to-day. Future work will need to explore if internal features were more relevant in our distance to the person average and associated activity in the left hemisphere.

### Identity processing in fMRI

When examining patterns of brain activity, face-identity representations have typically been found in the right FFA and the vATL and not in the left hemisphere (Kriegeskorte et al., 2007; Nestor et al., 2011; Verosky et al., 2013; Anzellotti et al. 2014; Axelrod & Yovel, 2015). Face stimuli in these studies, however, have had limited within-identity variability showing few or no changes in viewpoints, expressions, hair styles, etc. Low variability among faces simplifies the operations needed to match a particular face to its identity resulting in weak demands on within-identity face processing. Similarly, previous work used unfamiliar stimuli and tasks that prime exemplar-based representations, such as a one-back working memory task, resulting in weak activation of identity-related processing. In contrast, we used well-known faces associated with rich biographical information and actively engaged face identification on each trial by asking participants to specify the face with a button-press. Finally, prior work has focused solely on between-identity discrimination and found patterns of brain activity that differ by identity. We focused on within-identity discrimination and found brain regions displaying activity varied by how similar each photo looked like that person. Representations for within-identity of familiar faces may instead lie in the left hemisphere while representations for between-identity may lie in the right hemisphere.

We highlight that the use of behavioral and computational measures has a considerable impact on the localization of brain function. For instance, Jeong and Xu (2016) used a set of highly variable photos of famous faces, as in our study, to localize identity processing. In one analysis, the authors correlated patterns of brain activity with a behavioral measure (visual search speeds for finding a target actor among distractors). Unlike in our study, they found identity information in the superior intraparietal sulcus (IPS) but not in the FFA. Since visual search tasks engage short-term memory and activate the superior IPS (Todd & Marois, 2004; Xu & Chun, 2006), their behavioral measure likely tracked the aspect of person identity related to visual short-term memory and not identity processing more generally. Similarly, our use of the likeness ratings also operationalized a particular aspect of person identity, within-person identity, and not a global factor about identity. Consequently, while processing of face-identity is likely distributed across the brain, different regions will specialize in particular functions. This viewpoint encourages the application of different behavioral and computational measures to help elucidate the differing aspects in the neural representation of face and person identity.

### Exemplar models

One alternative to the prototype models used in the present study are exemplar models (Supplementary Results). In an exemplar approach, each face is encoded relative to exemplars of previously experienced faces (Valentine, 1991). The exemplar approach has been especially popular in visual category learning where they better account for behavioral and neuroimaging data when compared to prototype models (Mack et al., 2013; Richler & Palmeri, 2014). However, exemplar models have not fared as well in explaining face processing (Loffler et al., 2005; Chang & Tsao, 2017). However, when we compared the two approaches, we did not find a significant difference between prototype and exemplar models in explaining brain activity, a result similar to a prior simulation study (Ross et al., 2013). We only considered the distance to the average face and not the direction, which has been shown to be important in prior work (Leopold et al., 2006; Chang & Tsao, 2017), and should be considered for future work. We note that this distinction between the two models may not be as relevant for our work since exemplar models can represent prototypes implicitly (e.g., weighted activation of all exemplars can be equivalent to the average). Thus, the exact neural implementation (exemplar or prototype) is not as important for our conclusions as the particular representation, which we argue here is a prototype.

## Conclusion

Our results have significant implications for the neural mechanisms supporting familiar face and face-identity processing. First, by showing dissociable effects between hemispheres, our results argue that the left hemisphere makes unique contributions to face processing, in addition to the typical finding of right hemisphere dominance for faces. The left hemisphere represents within-person identity based on the person average, while the right hemisphere engages in face categorization based on the center average. The right hemisphere may represent the whole face-space comparing between people while the left hemisphere may represent the particular face-space of a familiar person. Second, by showing that weighting the distances to the person average fit brain activity the best, our results imply that person identity is represented by how someone looks on average scaled by the relative importance of each face feature. A feature is scaled based on how distinctive that feature is amongst the population. For instance, if an individual has large eyes, then this feature will be unimportant if everyone else also has large eyes. It is already known that face averaging is a type of summary representation that occurs automatically and in real-time (Haberman & Whitney, 2007; 2009). It may also be the case that such summary representations consider the relative importance of each feature for discrimination in real-time.

## Supplementary Materials

### Stimuli

**Supplementary Table 1.** Listed are the names of the target and distractor celebrities used in the experiment.

| Target Celebrities | Distractor Celebrities |
| --- | --- |
| Angelina Jolie | Anne Hathaway |
| Brad Pitt | Barack Obama |
| Jennifer Aniston | Bill Clinton |
| Johnny Depp | Dwayne Johnson |
| Julia Roberts | Ellen DeGeneres |
| Justin Timberlake | George Clooney |
| Oprah Winfrey | Halle Berry |
| Will Smith | Kate Winslet |
|  | Keira Knightley |
|  | Leonardo DiCaprio |
|  | Meryl Streep |
|  | Michael Jackson |
|  | Morgan Freeman |
|  | Sandra Bullock |
|  | Tina Fey |
|  | Tom Cruise |

### Computational measures

To examine the properties of OpenFace and to confirm its appropriateness for our analyses in face-space, we input frames from the 864 video dataset into the neural network. As there are many layers of processing, we selected for study an early, middle, and late layer in the neural network. We presumed that representations would become more abstract and face-specific in later layers. To characterize the representations of OpenFace, we compared the output at each layer (early, middle, later) with measures of identity, low-level features (pose/luminance), and high-level features (age/attractiveness). Briefly, we found that the 128-dimensional output of OpenFace are consistent with behavioral and neural models of face-space (Supplementary Figure 1). From early-to-late layers, face representations have (1) high accuracy in identifying faces (frames) of the same person (video), (2) develop pose and luminance invariance, and (3) develop associations with face traits (i.e., age and attractiveness. Thus, the 128-dimensional output would serve as appropriate representations of face-space.

**Supplementary Figure 1:**
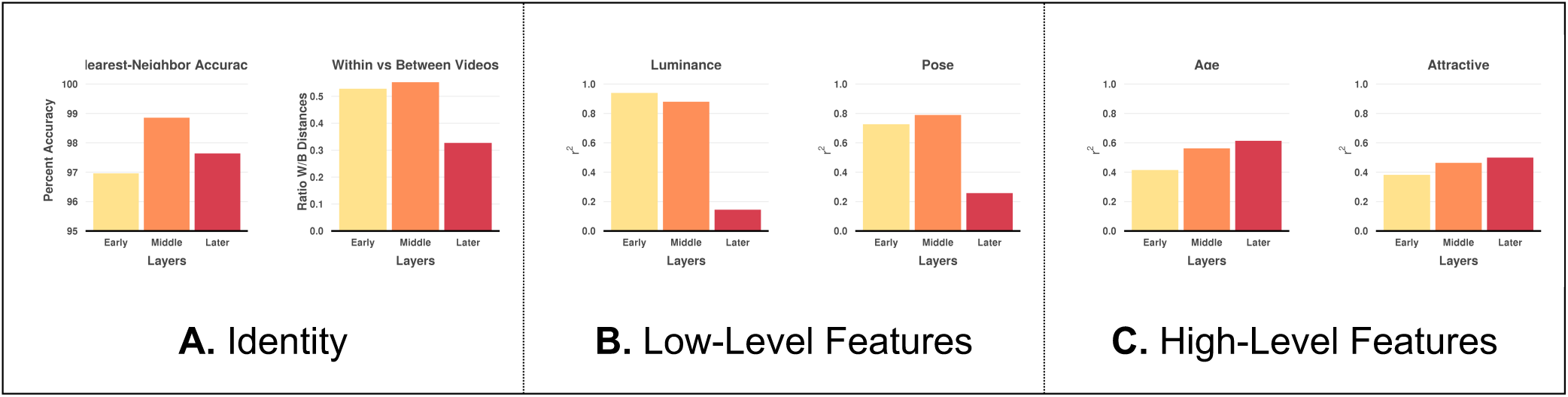
Features represented in the different layers of the neural network (x-axis) using faces from an 864 video dataset. For graphs with an *r^2^* in the y-axis, a ridge regression was used and the model fit is shown in terms of a cross-validated *r^2^*. (a) Face identification performance. On the left, percent accuracy that a frame in video is closest to other frame in video (y-axis) and on the right, the average distance within video (across frames) relative to between videos (y-axis). (b) Low-level features including predicting mean luminance on the left and pose on the right. (c) High-level features including predicting age on the left and attractiveness on the right.

### Exemplar measures

We compared our prototype-based measures (center average and person average) to equivalent exemplar-based measures. For both exemplar measures, we used the k-nearest-neighbors (kNN) approach and calculated the average distance to the nearest faces. We determined the number of nearest neighbors to sample by applying a kNN classifier with cross-validation to predict identity based on the 128-dimensional features in face-space, which resulted in an optimal *k* of 15. As a comparison to the center average, we calculated the average Euclidean distance between a photo and the nearest faces. As a comparison to the person average, we calculated the average Euclidean distance between a photo and that person’s nearest faces.

### Weighted distances to the person-average

Not all face features will be equally important for identity. To account for this, we developed another measure of distance to the person average that weights each feature by how important it is in identifying a person. We ran a multinomial ridge regression on the faces from our training set using the glmnet and caret R packages (Kuhn, 2008; Friedman et al., 2010). The 128 features for each face were used to predict the person’s identity (8 identities) with a 10-fold cross-validation repeated 10 times to estimate the penalization term (lambda). The model estimated separate feature weights for each identity that were then used to weight the Euclidean distance of each photo to that person’s average face.

### Preprocessing

Image preprocessing incorporated functions from AFNI (v 2014-10-23 OpenMP, http://afni.nimh.nih.gov/afni), FSL (v 5.0.7, http://www.fmrib.ox.ac.uk/fsl), and FreeSurfer (v 5.3.0, http://surfer.nmr.mgh.harvard.edu). The pipeline for functional images proceeded as follows: (1) the first 6 volumes (6s) were discarded to allow for MR equilibration, (2) motion correction was performed on the functional images using AFNI’s 3dvolreg, (3) skull stripping of the mean functional image was performed using FSL’s BET (Smith, 2002), (4) spatial smoothing with a 4-mm FWHM Gaussian mask was performed using FSL’s SUSAN (Smith & Brady, 1997), (5) high-pass filtering with a 0.01 Hz cut-off to remove low-frequency drift was conducted using FSL, as was mean-based global intensity normalization. Structural images were skull-stripped with a hybrid watershed/surface deformation procedure using FreeSurfer (Ségonne et al., 2004). The functional images were registered to the high-resolution structural images with boundary-based registration using FSL’s FLIRT. The structural images were in turn nonlinearly registered to the Montreal Neurological Institute’s MNI152 template (2 mm isotropic) using FSL’s FNIRT (Andersson et al., 2007) and this transform was then applied to the functional images in anatomical space.

### Regions-of-Interest (ROIs)

We used a probability map of face-selective responses that took the average of two localizers (face vs. house and face vs. scene) and represented the percentage of participants with a significant face-selective response (*p* < 0.05, one-tailed). To identify activation peaks, we smoothed the map by 2mm using AFNI’s 3dBlurInMask, and then used AFNI’s 3dExtrema to find the local maxima (minimum distance between peaks = 12 mm; probability threshold = 0.1). We selected those local maxima found in the fusiform gyrus and the lateral occipital cortex based on FreeSurfer’s cortical labeling of the MNI152 brain. Eight peak locations were found, including the left OFA (X = -44, Y = -82, Z = -10), the left FFA (X = -40, Y = -50, Z = - 18), the left aFus (X = -42, Y = -28, Z = -20), the left vATL (X = -34, Y = -6, Z = -34), the right OFA (X = 42, Y = -78, Z = -10), the right FFA (X = 42, Y = -50, Z = -20), the right aFus (X = 44, Y = -28, Z = -22), and the right vATL (X = 32, Y = -2, Z = -36). From the probabilistic map, we also found two peak locations in V1 to serve as control regions where we do not expect strong face-selective activity: the left V1 (X = -6, Y = -80, Z = 8) and the right V1 (X = 12, Y = - 76, Z = 12). For each peak, we created an ROI with a sphere of 4mm radius in MNI152 space (2 mm isotropic) using AFNI’s 3dUndump, which were then transformed into each participant’s functional space (2.5 mm isotropic).

### fMRI Data Analysis

Here we specify the analytic approach when using ROIs and voxelwise data, and then indicate the specific models that were run.

#### Analytic approach

##### Regions of Interest

We took the mean time-series within each ROI from the probabilistic atlas. At the subject-level, we applied multiple linear regression on the ROI time-series to estimate the beta-coefficients for each regressor and contrast. The coefficients for a given regressor or contrast were then used at the group-level in a t-test to calculate the significance across participants.

##### Voxelwise

Whole-brain voxelwise regression analyses were performed at the subject-level using AFNI’s 3dDeconvolve and 3dREMLfit functions, which provide temporal pre-whitening via an autoregressive model. Following individual analyses, whole-brain group analyses were performed using AFNI’s 3dMEMA (Chen et al., 2012), which incorporates individual beta precision estimates into group effects using a mixed-effects meta-analytic approach. Clusters were defined as contiguous sets of voxels with Z > 1.96 and then thresholded using Gaussian random field theory (cluster probability *p* < 0.05) to correct for multiple comparisons (Worsley et al., 1996).

### Regression models

#### Brain activity for faces

We sought to find regions with significant activity for faces versus scenes. The linear model consisted of two explanatory variables for trials with a target face and trials with a distractor face convolved with a double-gamma function (AFNI’s SPMG1). Regressors of no interest included baseline effects of each run and six head movement parameters. Additional regressors were given for target stimuli, which were convolved with a double-gamma function, included trials with incorrect responses, trials with no responses, and the amplitude-modulated response time to account for the effect of button presses. Finally, a contrast was defined to identify regions that showed increased activity when viewing both target and distractor faces. Since scene trials served as the baseline and weren’t explicitly modeled, a face contrast was implicitly face > scene.

#### Brain activity for face familiarity

We took the mean response time for each identity across photos for representing the level of familiarity of a participant with an identity. The linear model consisted of one explanatory variable where each target trial indicates the level of familiarity with that identity (mean response time) convolved with a double-gamma function. The rest of the model was the same as the prior model for face activity. This analysis was only done for the ROIs.

#### Associations with the center average and the person average

Given that the stimuli were highly familiar to participants, we wanted to assess the importance of face averages in the neural representation of familiar faces. This model consisted of two explanatory variables. For each target trial, we modeled the distance of the photo to the center average and to the person average, and convolved both variables with a double-gamma function. The rest of the model was the same as the previous analysis except we did not include response time due to potential collinearities with our face average measures. In addition, to account for any between-identity effects, we also included regressors of no interest that modeled activity on target trials for each person identity. To assess the significance of the most discriminable face features in representing the person average, we repeated the analysis and replaced the distance to the person average regressor with that of the weighted distance to the person average.

#### Associations with k-nearest-neighbors (kNN)

To test an exemplar model, we used kNN. We repeated the analysis for the center average and the weighted person average (our prototype model) except replaced those prototype measures with the overall or person-specific kNN measures (our exemplar model). We then compared the outputs from the prototype and exemplar model to see which provided the best fit for the data.

#### Associations with likeness ratings and response times

We compared our behavioral measures with brain activity. Could we identify brain regions that varied with how much a photo looked like a person (likeness ratings) and how quickly the person in the photo was identified (reaction time)? This model included two explanatory variables convolved with a double-gamma function, one for likeness ratings and another for response times on each target trial. Regressors of no interest were the same as the previous analysis (associations with the center and person average).

#### Mediation

We ran a mediation analysis (only on our ROIs) to find if our behavioral measures would mediate the effect of distance to the center/person average on brain activity. We used the R package mediation with a quasi-Bayesian approximation (100 simulations) to measure the average causal mediation effect (indirect effect) and the proportion of the effect mediated.

## Supplementary Results

### Behavioral Results

**Supplementary Table 2.** Listed are the mean and standard deviations of behavioral and computational measures (columns) for each celebrity’s photos (rows).

| Celebrity | Likeness Ratings | Response Time | Distance to Center Average | Distance to Person Average |
| --- | --- | --- | --- | --- |
| *Angelina Jolie* | 4.24 ± 0.87 | 0.88 ± 0.06 | 0.98 ± 0.02 | 0.57 ± 0.12 |
| *Brad Pitt* | 4.03 ± 0.54 | 0.85 ± 0.06 | 0.95 ± 0.03 | 0.50 ± 0.10 |
| *Jennifer Aniston* | 4.09 ± 0.86 | 0.75 ± 0.05 | 0.98 ± 0.04 | 0.54 ± 0.13 |
| *Johnny Depp* | 3.93 ± 0.58 | 0.84 ± 0.06 | 0.92 ± 0.03 | 0.66 ± 0.11 |
| *Julia Roberts* | 4.33 ± 0.68 | 0.90 ± 0.04 | 0.95 ± 0.03 | 0.47 ± 0.12 |
| *Justin*<br>*Timberlake* | 3.69 ± 0.38 | 0.79 ± 0.04 | 0.94 ± 0.03 | 0.51 ± 0.15 |
| *Oprah Winfrey* | 3.86 ± 0.35 | 0.75 ± 0.05 | 1.04 ± 0.02 | 0.30 ± 0.08 |
| *Will Smith* | 4.13 ± 0.85 | 0.78 ± 0.06 | 1.05 ± 0.03 | 0.46 ± 0.12 |

### Face Activity

We contrasted brain activity for faces versus scenes in ROIs (Supplementary Figure 2). Brain activity significantly differed between the hemispheres (F_1,99_=4.3, *p* < 0.05) and was lateralized to the right hemisphere. In addition, the ROIs significantly differed in their level of brain activity (F_9,99_=10.4, *p* < 0.05) with activity peaking in the right FFA ROI. Almost every region in the right hemisphere was significant except V1 (significant regions included the OFA, FFA, aFus, vATL), while in the left hemisphere, only the OFA and the FFA were significant (all *ps* < 0.05). It is relevant that the response to faces > scenes was not significant in V1, since this was a control ROI and a face-selective response was not expected.

**Supplementary Figure 2:**
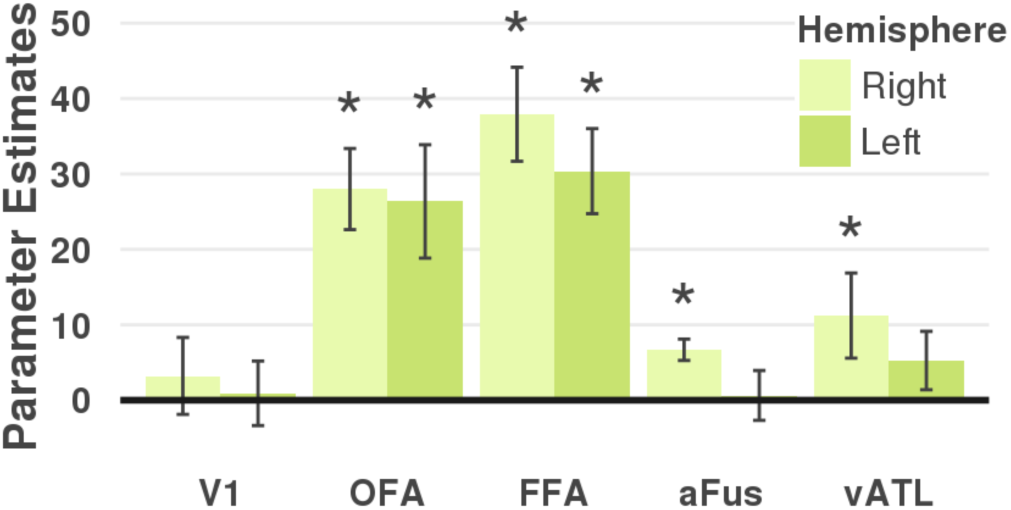
Bar plots showing brain activity of seeing faces > scenes (beta parameter estimates; y-axis) for each ROI (x-axis). Lighter color bars are for ROIs in the right hemisphere while darker color bars are for the left hemisphere. An asterisk above a bar indicates significance (*p* < 0.05, two-tailed).

**Supplementary Table 3a:**
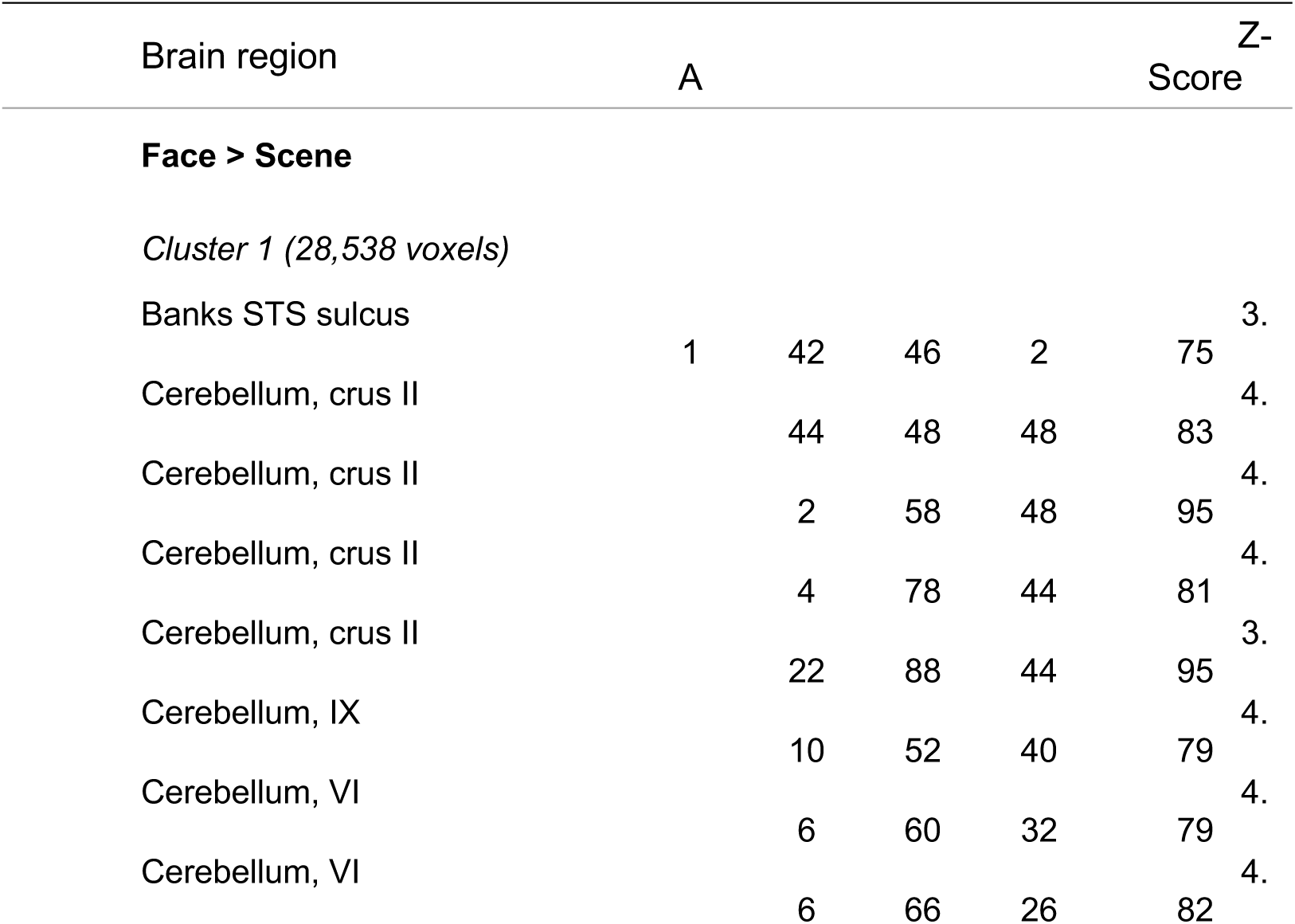
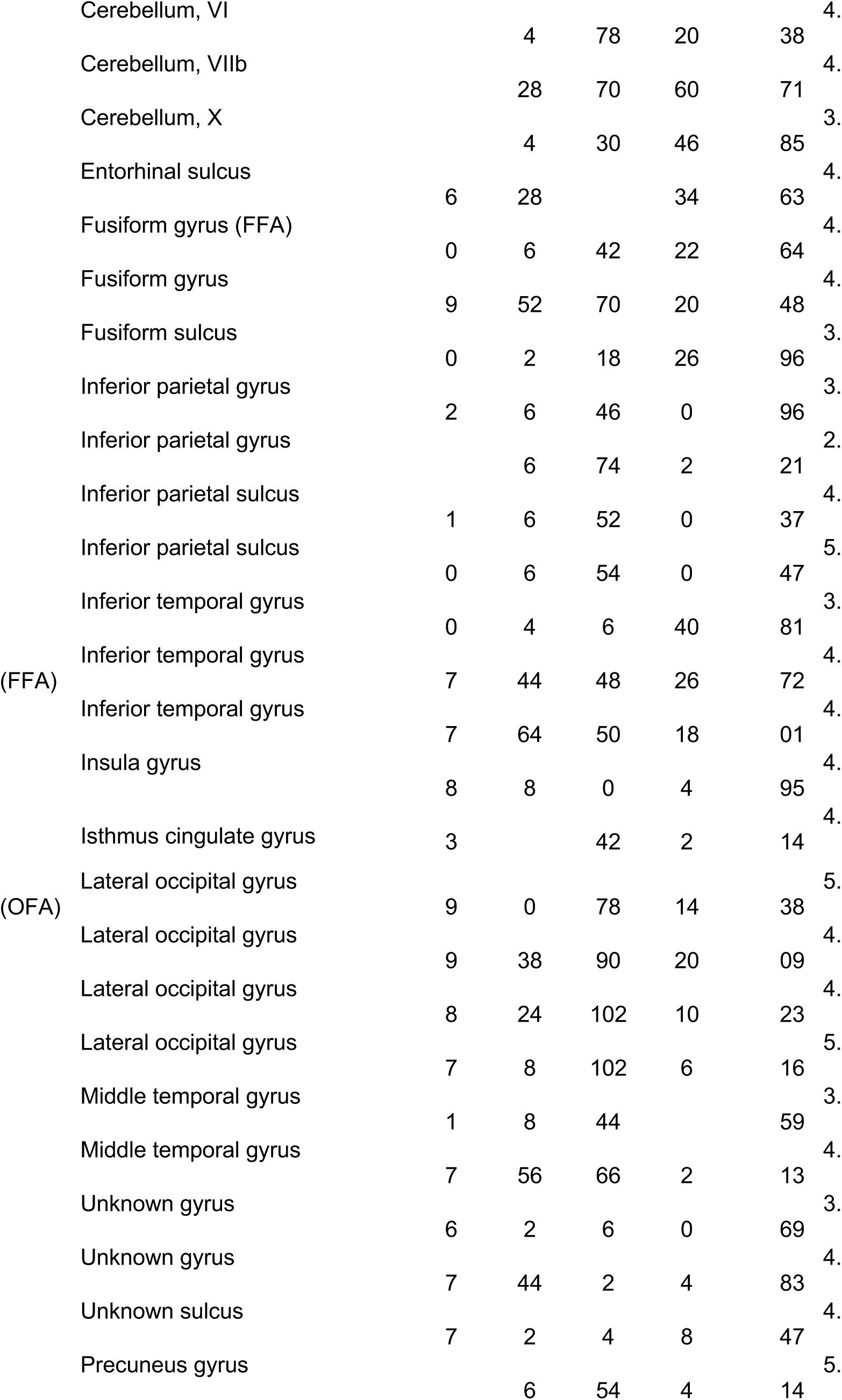
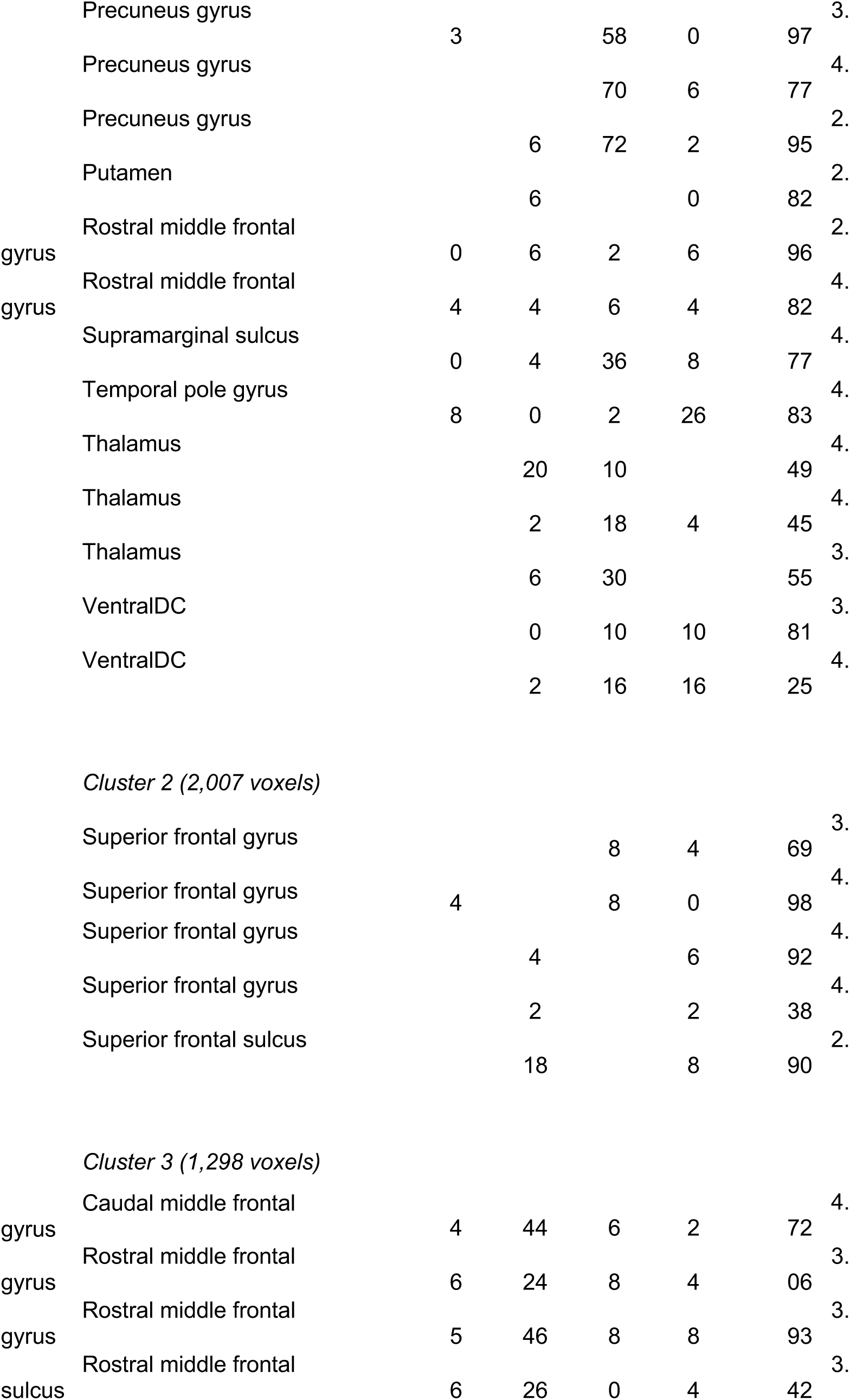
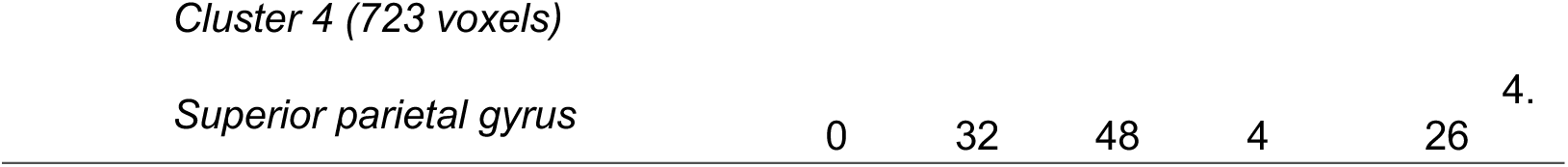
Listed are the peaks of significant ***brain activity for faces > scenes***. Peaks were found using AFNI’s 3dExtrema by setting a voxel-level threshold of Z > 1.96 (cluster corrected, *p* < 0.05) and a minimum distance of 20 mm between peaks. Abbreviations: H, hemisphere; B, bilateral; R, right; L, left; BA = Brodmann Area.

**Supplementary Table 3b:**
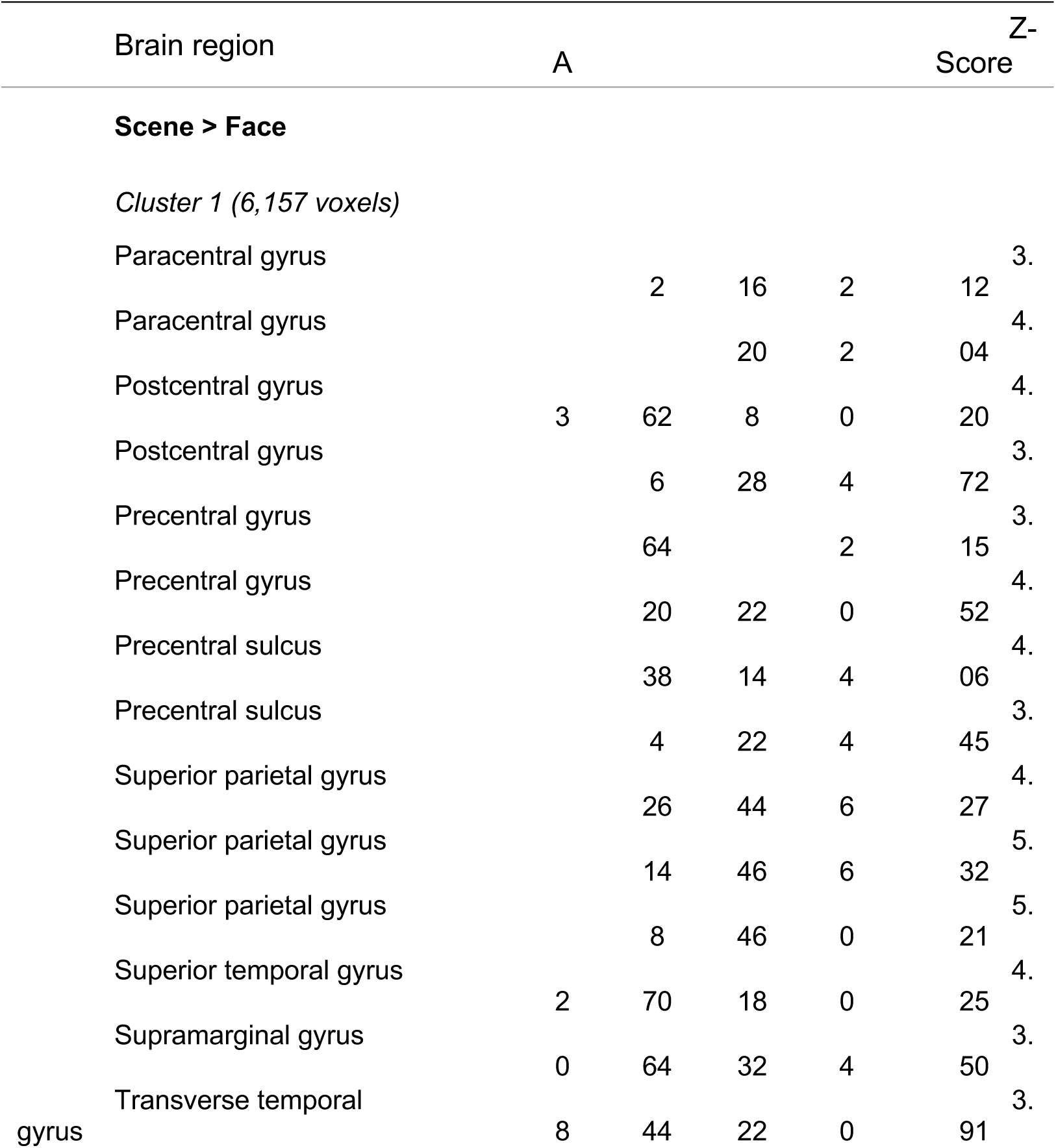
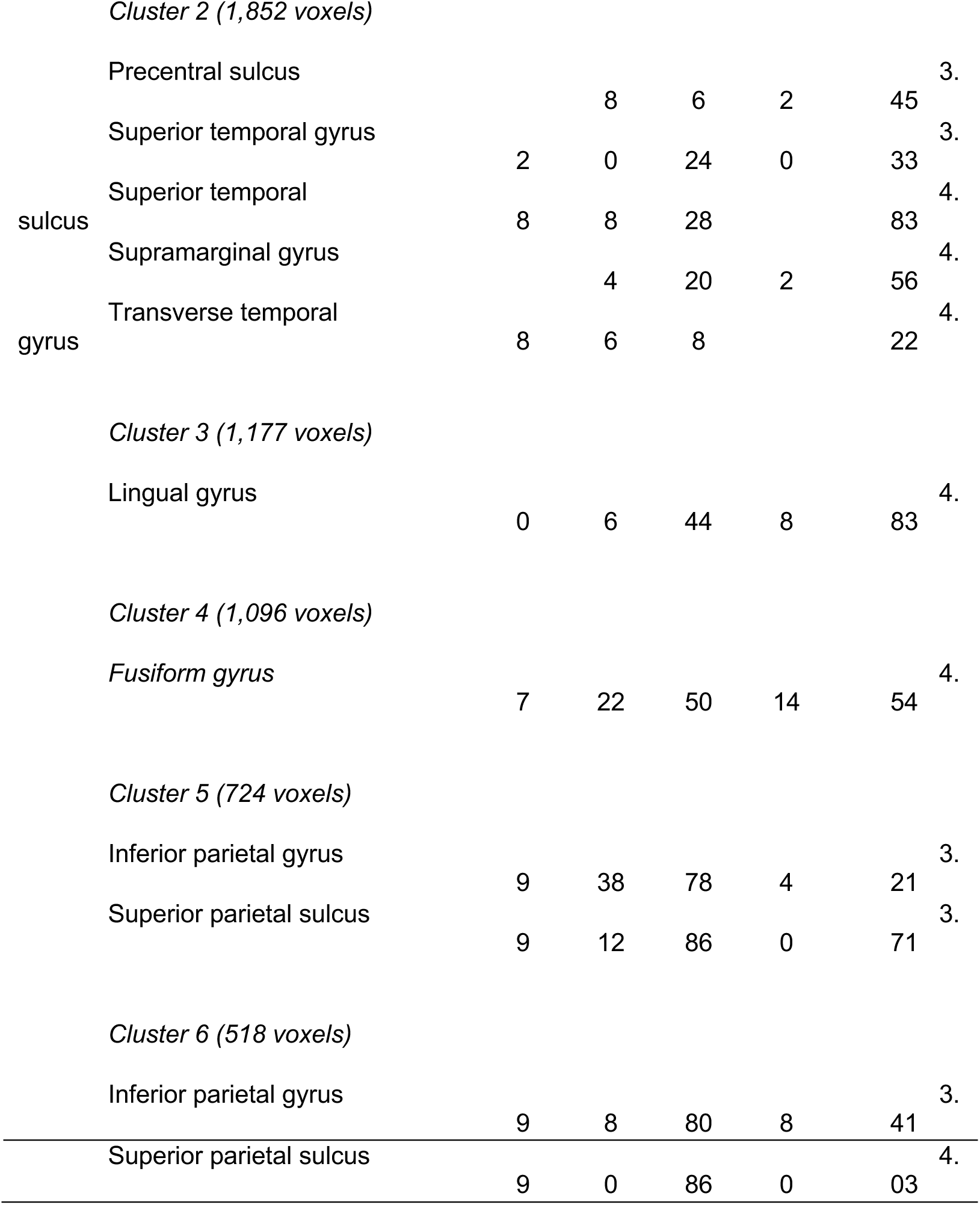
Listed are the peaks of significant ***brain activity for scenes > faces***. Peaks were found using AFNI’s 3dExtrema by setting a voxel-level threshold of Z > 1.96 (cluster corrected, *p* < 0.05) and a minimum distance of 20 mm between peaks. Abbreviations: H, hemisphere; B, bilateral; R, right; L, left; BA = Brodmann Area.

### Distances to the Center Average and Person Average

We examined the relationship between brain activity and distances to the center and person averages in our ROIs. First, when explaining variation in brain activity with the distance to this center average, we did not find any main effect of hemisphere or ROI (all *ps* > 0.05). However, we found significant associations in the right FFA (t_5_=-3.1, *p* < 0.05) and right aFus (t_5_=-2.2, *p* < 0.05); photos that were more face-like (i.e., closer to the center average) tended to activate the right FFA and the right aFus. Second, when explaining variation in brain activity with the distance to this person average, we found a significant main effect of ROI (F_9,99_=2.0, *p* < 0.05) but not of hemisphere. For each individual ROI, we found significant positive associations (*p* < 0.05) in the right OFA (t_5_=3.5), the right FFA (t_5_=2.0), the left FFA (t_5_=3.2), and the left aFus (t_5_=2.7); photos that looked less like that person (i.e., further from the person average) tended to activate those regions. Of those regions (OFA, FFA, aFus), only the effect in the left aFus was significantly greater than the associated right hemisphere aFus region (t_5_=3.0, *p* < 0.05).

**Figure 11:**
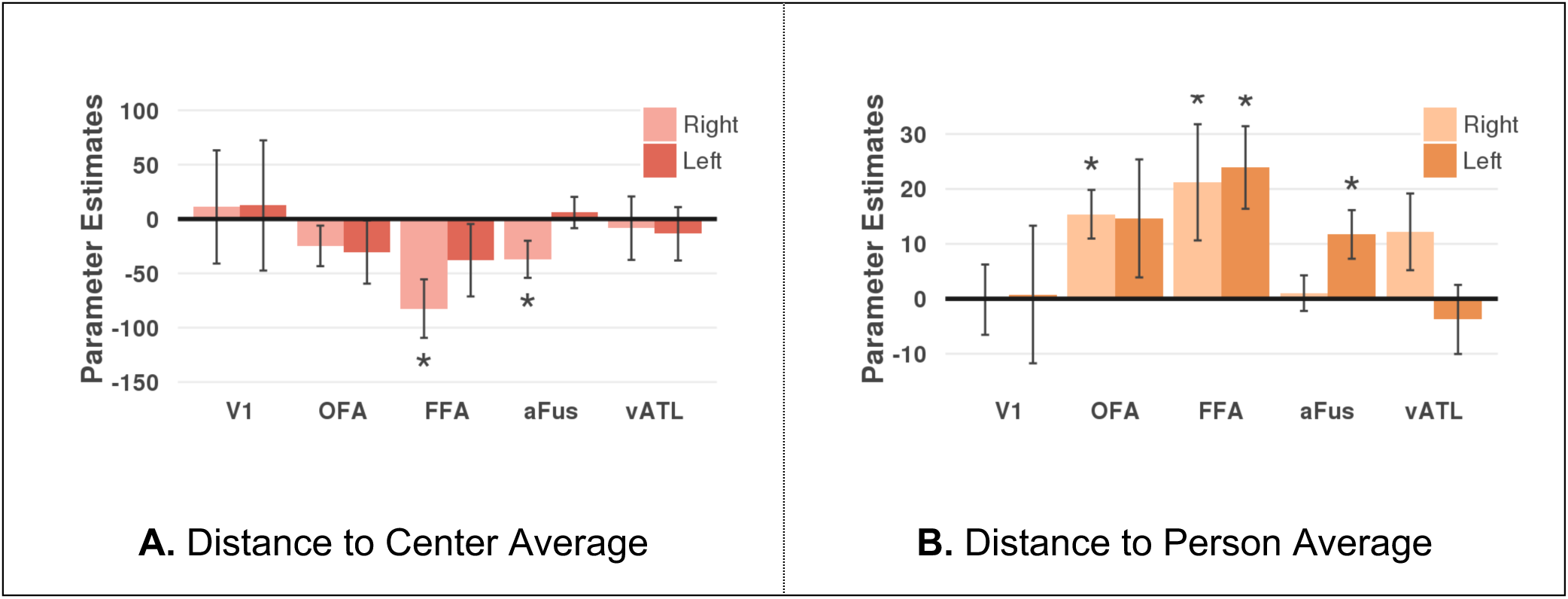
Significance of brain activity (y-axis) associated with distances of each photo to the center average (left, red) and the person average (right, orange) is shown for each ROI (x-axis). Negative associations indicate that brain signal increases for photos closer to an average face while positive associations indicate that brain signal increases the further a photo is to an average face. Lighter color bars are for ROIs in the right hemisphere while darker color bars are for the left hemisphere. An asterisk above a bar indicates significance (*p* < 0.05, two-tailed).

### Weighted features by discriminability of identity

In the previous analysis, each feature was one of the 128 dimensions in face-space and with equal weight when calculating distances to person averages. Since some features might be more discriminative of a person’s identity, we weighted each feature based on its importance in classifying person identity with different weights for each person. Across all ROIs except V1, the weighted versus unweighted distances showed a stronger association with brain activity (Supplementary Figure 3). To directly compare the benefit of the weighted over the unweighted distances, we added the weighted distances as a regressor to a model with the unweighted. We found a significant effect of weighted distances over and above unweighted distances in the right OFA and the aFus, as well as the left OFA, FFA, and aFus. When we instead added the unweighted distances to a model with the weighted distances, no region showed a significant effect on brain activity for the unweighted distances (all *ps* > 0.05).

**Supplementary Figure 3:**
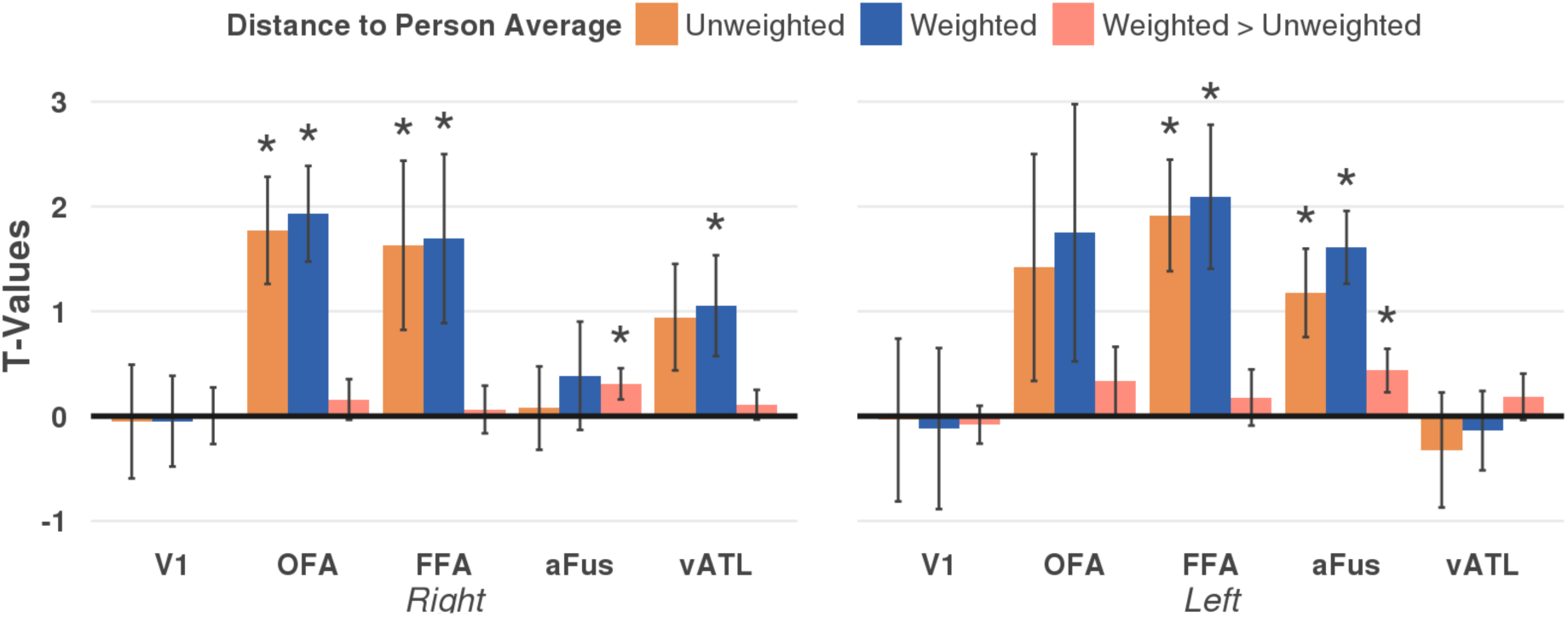
Bar graphs showing the effect of weighting the distances on explaining brain activity as T-values (y-axis) for the different ROIs (x-axis). Distances of each photo to that person’s average face were calculated with each feature having the same weight (unweighted, orange) or each feature weighted based on the coefficients of a multinomial ridge regression where features that were more important in discriminating an identity had higher values (weighted, blue). The ridge regression used the 128 features of the training data to classify the 8 identities and produce feature weights that were unique to each identity; then these weights were applied to the test data. The pink bar shows the additional effect of adding the weighted distances to a model with the unweighted distances (weighted > unweighted).

**Supplementary Table 4:**
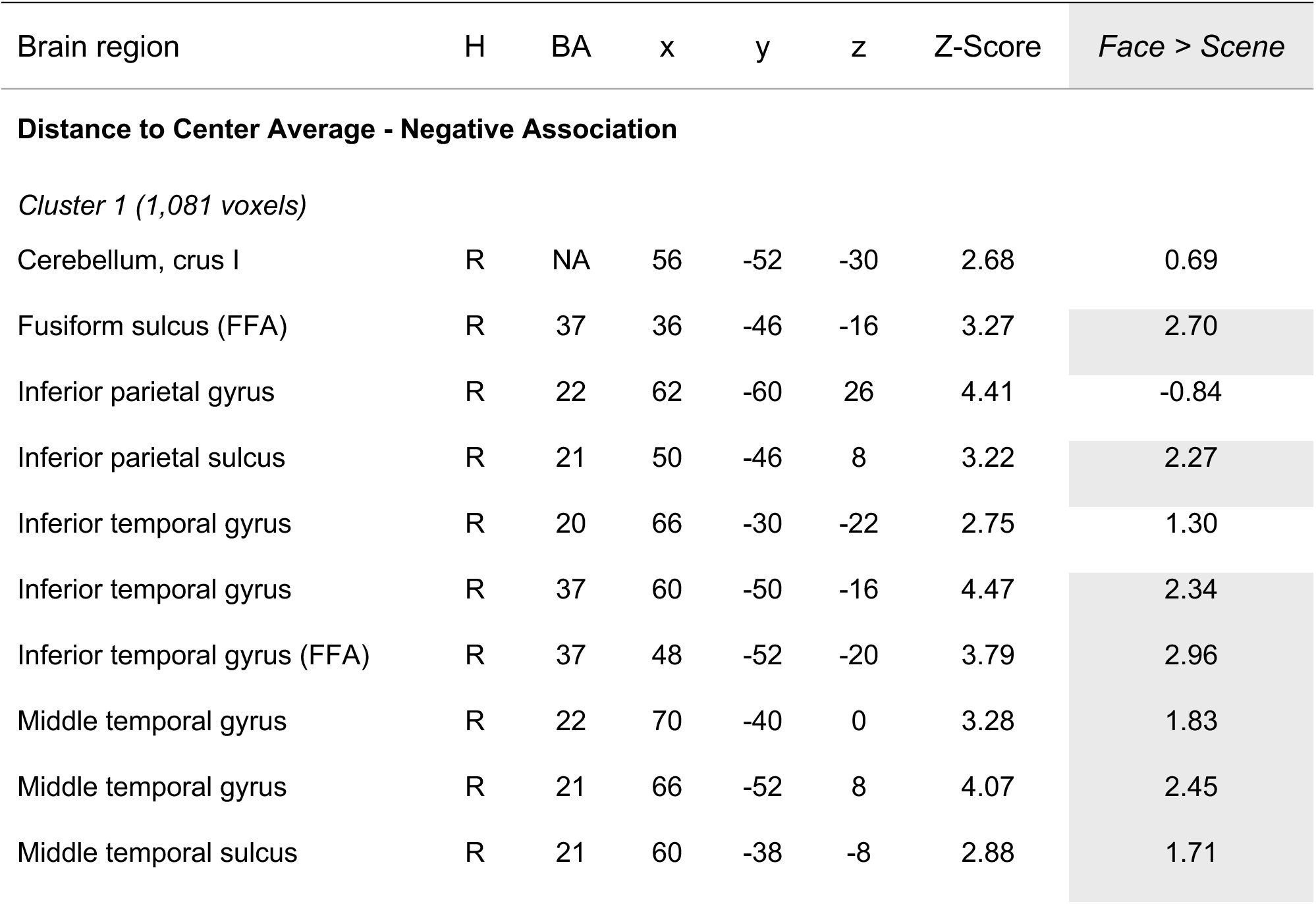
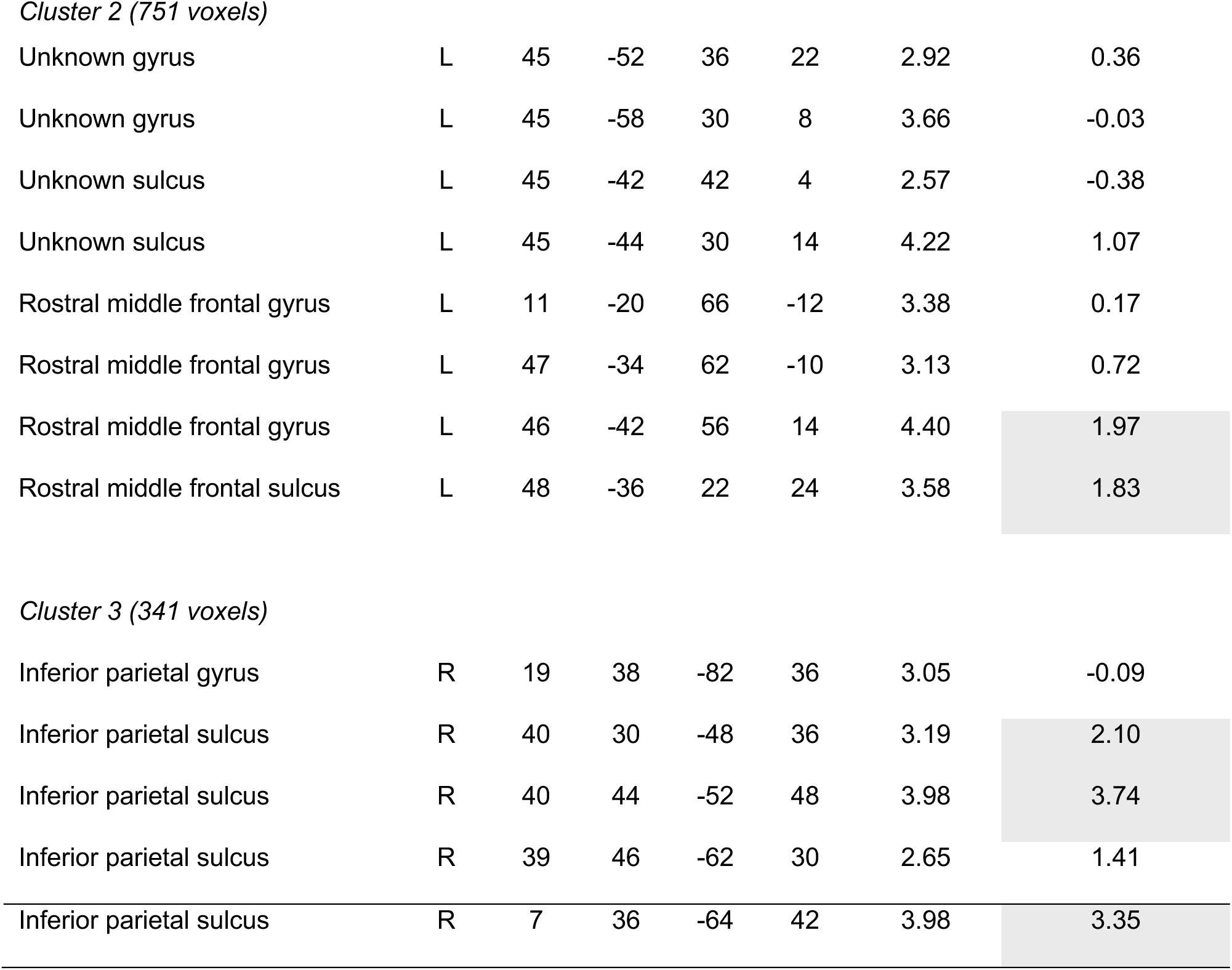
Listed are the peaks of significant ***brain activity negatively associated with the distance of each photo to the center average***. Peaks were found using AFNI’s 3dExtrema by setting a voxel-level threshold of Z > 1.96 (cluster corrected, *p* < 0.05) and a minimum distance of 12 mm between peaks. The last column provides the Z-Score for the Face > Scene contrast. Abbreviations: H, hemisphere; B, bilateral; R, right; L, left; BA = Brodmann Area.

| Brain region | H | BA | x | y | z | Z-Score | Face > Scene |
| --- | --- | --- | --- | --- | --- | --- | --- |
| **Distance to Center Average - Negative Association** |  |  |  |  |  |  |  |
| *Cluster 1 (1,081 voxels)* |  |  |  |  |  |  |  |
| Cerebellum, crus I | R | NA | 56 | -52 | -30 | 2.68 | 0.69 |
| Fusiform sulcus (FFA) | R | 37 | 36 | -46 | -16 | 3.27 | 2.70 |
| Inferior parietal gyrus | R | 22 | 62 | -60 | 26 | 4.41 | -0.84 |
| Inferior parietal sulcus | R | 21 | 50 | -46 | 8 | 3.22 | 2.27 |
| Inferior temporal gyrus | R | 20 | 66 | -30 | -22 | 2.75 | 1.30 |
| Inferior temporal gyrus | R | 37 | 60 | -50 | -16 | 4.47 | 2.34 |
| Inferior temporal gyrus (FFA) | R | 37 | 48 | -52 | -20 | 3.79 | 2.96 |
| Middle temporal gyrus | R | 22 | 70 | -40 | 0 | 3.28 | 1.83 |
| Middle temporal gyrus | R | 21 | 66 | -52 | 8 | 4.07 | 2.45 |
| Middle temporal sulcus | R | 21 | 60 | -38 | -8 | 2.88 | 1.71 |
| *Cluster 2 (751 voxels)* |  |  |  |  |  |  |  |
| Unknown gyrus | L | 45 | -52 | 36 | 22 | 2.92 | 0.36 |
| Unknown gyrus | L | 45 | -58 | 30 | 8 | 3.66 | -0.03 |
| Unknown sulcus | L | 45 | -42 | 42 | 4 | 2.57 | -0.38 |
| Unknown sulcus | L | 45 | -44 | 30 | 14 | 4.22 | 1.07 |
| Rostral middle frontal gyrus | L | 11 | -20 | 66 | -12 | 3.38 | 0.17 |
| Rostral middle frontal gyrus | L | 47 | -34 | 62 | -10 | 3.13 | 0.72 |
| Rostral middle frontal gyrus | L | 46 | -42 | 56 | 14 | 4.40 | 1.97 |
| Rostral middle frontal sulcus | L | 48 | -36 | 22 | 24 | 3.58 | 1.83 |
| *Cluster 3 (341 voxels)* |  |  |  |  |  |  |  |
| Inferior parietal gyrus | R | 19 | 38 | -82 | 36 | 3.05 | -0.09 |
| Inferior parietal sulcus | R | 40 | 30 | -48 | 36 | 3.19 | 2.10 |
| Inferior parietal sulcus | R | 40 | 44 | -52 | 48 | 3.98 | 3.74 |
| Inferior parietal sulcus | R | 39 | 46 | -62 | 30 | 2.65 | 1.41 |
| Inferior parietal sulcus | R | 7 | 36 | -64 | 42 | 3.98 | 3.35 |

**Supplementary Table 5a:** Listed are the peaks of significant ***brain activity positively associated with the distance of each photo to the person average***. Peaks were found using AFNI’s 3dExtrema by setting a voxel-level threshold of Z > 1.96 (cluster corrected, *p* < 0.05) and a minimum distance of 12 mm between peaks. The last column provides the Z-Score for the Face > Scene contrast. Abbreviations: H, hemisphere; B, bilateral; R, right; L, left; BA = Brodmann Area.

| Brain region | H | BA | x | y | z | Z-Score | Face > Scene |
| --- | --- | --- | --- | --- | --- | --- | --- |

| Distance to Person Average - Positive Associations |  |  |  |  |  |  |  |
| --- | --- | --- | --- | --- | --- | --- | --- |
| *Cluster 1 (1,360 voxels)* |  |  |  |  |  |  |  |
| Cerebellum, crus I | L |  | -24 | -86 | -22 | 3.40 | 2.20 |
| Fusiform gyrus | L | 20 | -34 | -26 | -28 | 2.91 | 0.44 |
| Fusiform gyrus (FFA) | L | 37 | -36 | -54 | -14 | 3.81 | 2.25 |
| Fusiform gyrus | L | 37 | -46 | -64 | -10 | 2.25 | 1.51 |
| Fusiform gyrus | L | 19 | -34 | -70 | -14 | 3.78 | 2.61 |
| Inferior temporal gyrus | L | 20 | -44 | -40 | -20 | 4.25 | 2.60 |
| Inferior temporal gyrus | L | 37 | -50 | -54 | -16 | 3.26 | 1.92 |
| Inferior temporal sulcus (aFus) | L | 20 | -46 | -28 | -22 | 3.73 | 1.66 |
| Lateral occipital gyrus | L | 19 | -44 | -82 | 14 | 3.39 | -0.90 |
| Lateral occipital gyrus | L | 18 | -36 | -90 | 6 | 4.40 | 0.45 |
| Lateral occipital sulcus | L | 18 | -16 | -84 | -12 | 3.02 | -0.50 |
| Lateral occipital sulcus | L | 19 | -34 | -90 | -10 | 2.99 | 2.30 |
| *Cluster 2 (530 voxels)* |  |  |  |  |  |  |  |
| Fusiform sulcus | R | 37 | 44 | -66 | -10 | 2.15 | 1.75 |
| Lateral occipital gyrus (OFA) | R | 19 | 48 | -80 | -10 | 2.27 | 3.10 |
| Lateral occipital gyrus | R | 19 | 40 | -90 | -4 | 3.60 | 2.73 |
| Lateral occipital gyrus | R | 18 | 24 | -96 | -8 | 2.54 | 2.55 |
| Lateral occipital sulcus | R | 19 | 42 | -76 | 4 | 2.79 | 0.02 |
| Lateral occipital sulcus (OFA) | R | 19 | 36 | -80 | -10 | 3.48 | 2.03 |
| Lingual gyrus | R | 18 | 14 | -84 | -10 | 3.04 | 2.41 |

**Supplementary Table 5b:**
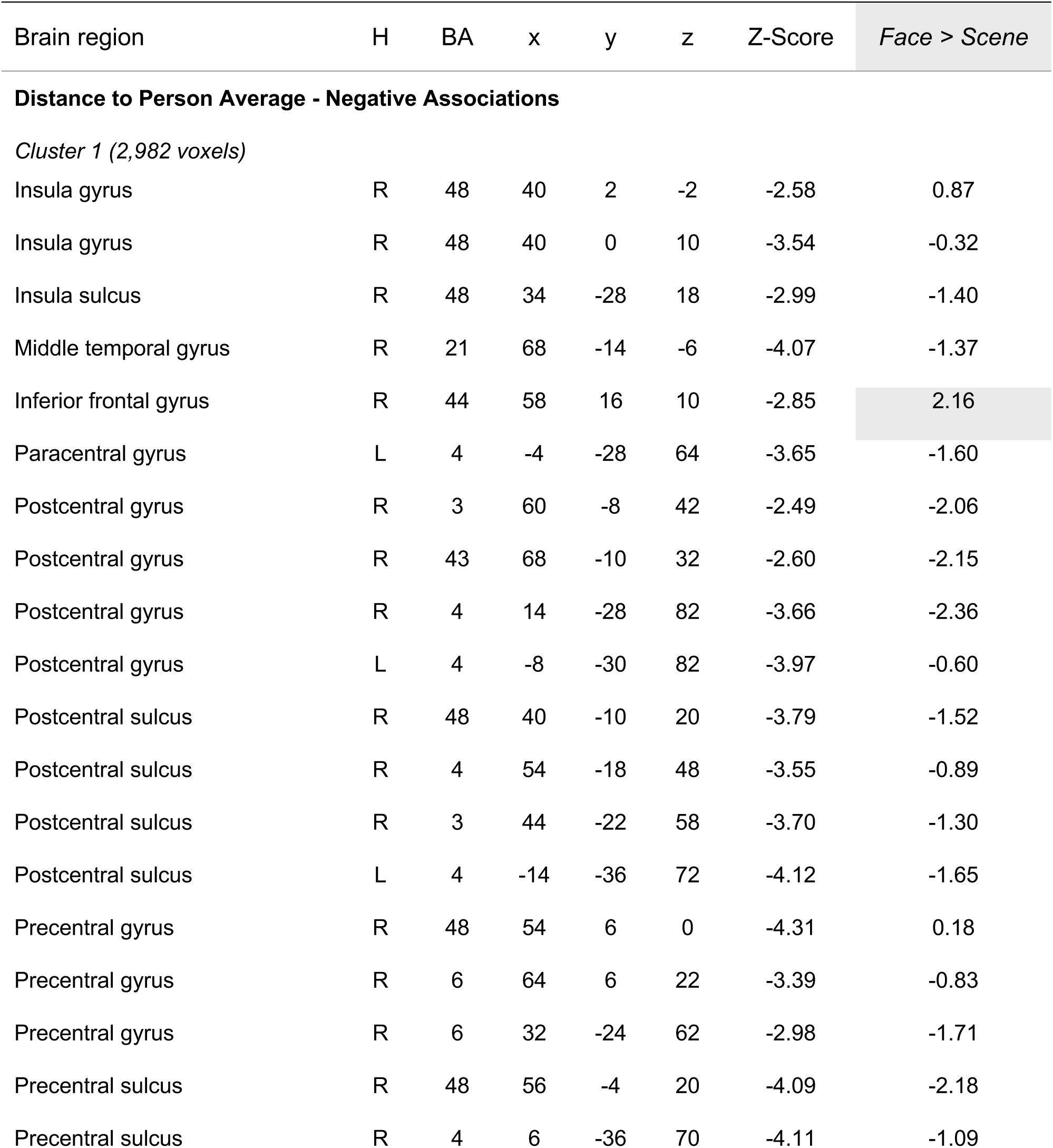
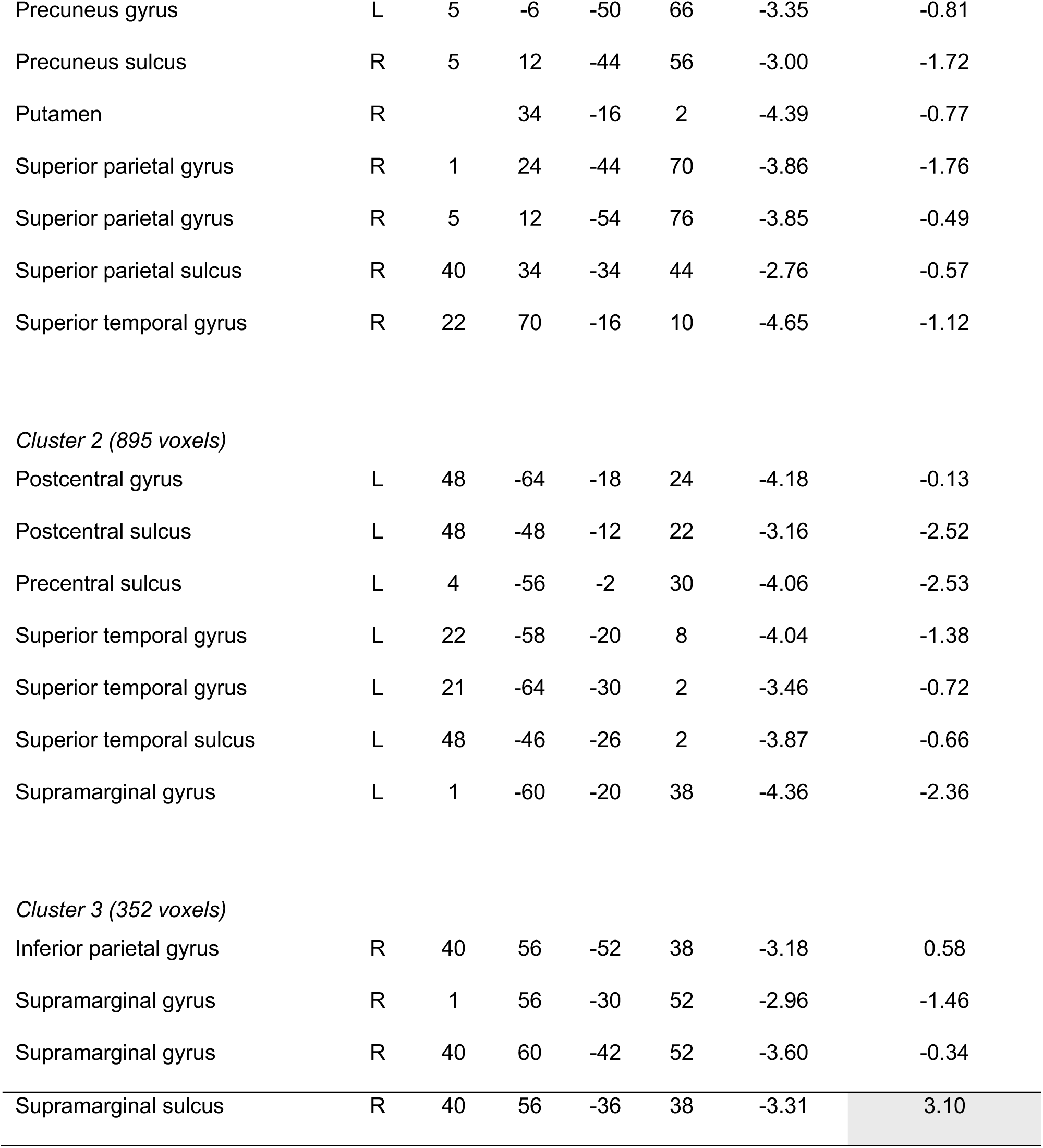
Listed are the peaks of significant ***brain activity negatively associated with the distance of each photo to the person average***. Peaks were found using AFNI’s 3dExtrema by setting a voxel-level threshold of Z > 1.96 (cluster corrected, *p* < 0.05) and a minimum distance of 12 mm between peaks. The last column provides the Z-Score for the Face > Scene contrast. Abbreviations: H, hemisphere; B, bilateral; R, right; L, left; BA = Brodmann Area.

| Brain region | H | BA | x | y | z | Z-Score | Face > Scene |
| --- | --- | --- | --- | --- | --- | --- | --- |
| **Distance to Person Average - Negative Associations** |  |  |  |  |  |  |  |
| *Cluster 1 (2,982 voxels)* |  |  |  |  |  |  |  |
| Insula gyrus | R | 48 | 40 | 2 | -2 | -2.58 | 0.87 |
| Insula gyrus | R | 48 | 40 | 0 | 10 | -3.54 | -0.32 |
| Insula sulcus | R | 48 | 34 | -28 | 18 | -2.99 | -1.40 |
| Middle temporal gyrus | R | 21 | 68 | -14 | -6 | -4.07 | -1.37 |
| Inferior frontal gyrus | R | 44 | 58 | 16 | 10 | -2.85 | 2.16 |
| Paracentral gyrus | L | 4 | -4 | -28 | 64 | -3.65 | -1.60 |
| Postcentral gyrus | R | 3 | 60 | -8 | 42 | -2.49 | -2.06 |
| Postcentral gyrus | R | 43 | 68 | -10 | 32 | -2.60 | -2.15 |
| Postcentral gyrus | R | 4 | 14 | -28 | 82 | -3.66 | -2.36 |
| Postcentral gyrus | L | 4 | -8 | -30 | 82 | -3.97 | -0.60 |
| Postcentral sulcus | R | 48 | 40 | -10 | 20 | -3.79 | -1.52 |
| Postcentral sulcus | R | 4 | 54 | -18 | 48 | -3.55 | -0.89 |
| Postcentral sulcus | R | 3 | 44 | -22 | 58 | -3.70 | -1.30 |
| Postcentral sulcus | L | 4 | -14 | -36 | 72 | -4.12 | -1.65 |
| Precentral gyrus | R | 48 | 54 | 6 | 0 | -4.31 | 0.18 |
| Precentral gyrus | R | 6 | 64 | 6 | 22 | -3.39 | -0.83 |
| Precentral gyrus | R | 6 | 32 | -24 | 62 | -2.98 | -1.71 |
| Precentral sulcus | R | 48 | 56 | -4 | 20 | -4.09 | -2.18 |
| Precentral sulcus | R | 4 | 6 | -36 | 70 | -4.11 | -1.09 |
| Precuneus gyrus | L | 5 | -6 | -50 | 66 | -3.35 | -0.81 |
| Precuneus sulcus | R | 5 | 12 | -44 | 56 | -3.00 | -1.72 |
| Putamen | R |  | 34 | -16 | 2 | -4.39 | -0.77 |
| Superior parietal gyrus | R | 1 | 24 | -44 | 70 | -3.86 | -1.76 |
| Superior parietal gyrus | R | 5 | 12 | -54 | 76 | -3.85 | -0.49 |
| Superior parietal sulcus | R | 40 | 34 | -34 | 44 | -2.76 | -0.57 |
| Superior temporal gyrus | R | 22 | 70 | -16 | 10 | -4.65 | -1.12 |
| *Cluster 2 (895 voxels)* |  |  |  |  |  |  |  |
| Postcentral gyrus | L | 48 | -64 | -18 | 24 | -4.18 | -0.13 |
| Postcentral sulcus | L | 48 | -48 | -12 | 22 | -3.16 | -2.52 |
| Precentral sulcus | L | 4 | -56 | -2 | 30 | -4.06 | -2.53 |
| Superior temporal gyrus | L | 22 | -58 | -20 | 8 | -4.04 | -1.38 |
| Superior temporal gyrus | L | 21 | -64 | -30 | 2 | -3.46 | -0.72 |
| Superior temporal sulcus | L | 48 | -46 | -26 | 2 | -3.87 | -0.66 |
| Supramarginal gyrus | L | 1 | -60 | -20 | 38 | -4.36 | -2.36 |
| *Cluster 3 (352 voxels)* |  |  |  |  |  |  |  |
| Inferior parietal gyrus | R | 40 | 56 | -52 | 38 | -3.18 | 0.58 |
| Supramarginal gyrus | R | 1 | 56 | -30 | 52 | -2.96 | -1.46 |
| Supramarginal gyrus | R | 40 | 60 | -42 | 52 | -3.60 | -0.34 |
| Supramarginal sulcus | R | 40 | 56 | -36 | 38 | -3.31 | 3.10 |

### Mediation Effects of Likeness Ratings

We examined if the effect of our face-space measures onto brain activity might be explained by our behavioral measure of likeness ratings. Thus, we performed a mediation analysis. While likeness ratings significantly mediated the effect of the center average on brain activity in almost every region, the proportion of the effect mediated tended to be small, less than 5% (Supplementary Figure 4). Likeness ratings strongly mediated the proportion of the person average effect. For the center average, the proportion mediated by likeness ratings was largest in the right FFA. While for the person average, the proportion mediated by likeness ratings was largest in the left aFus.

**Supplementary Figure 4:**
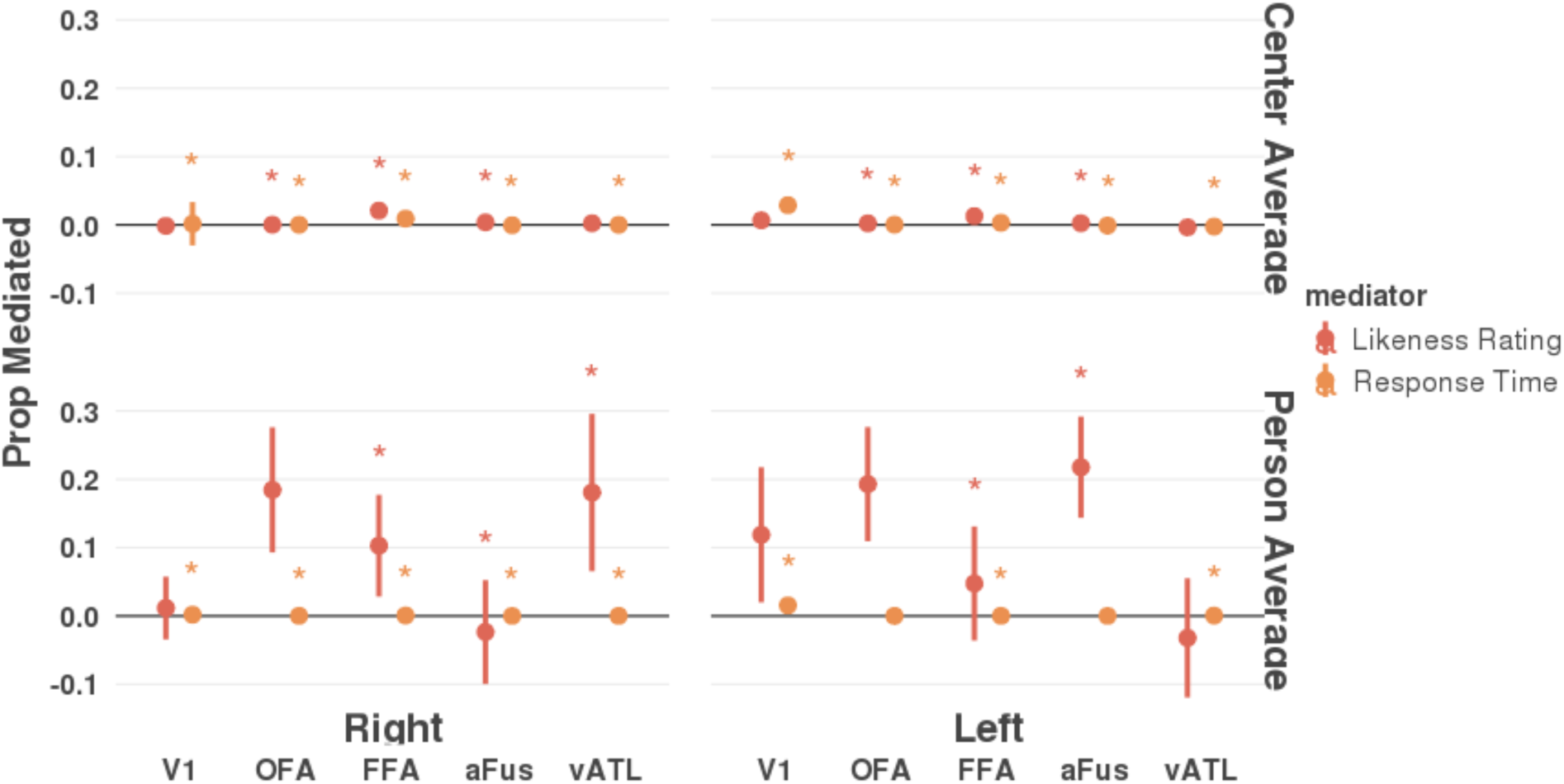
Plot shows results of a mediation analysis. The direct model tested how well brain activity in different ROIs (x-axis) could be explained by the distances between each photo and (top) the person average or (bottom) the center average (average of all faces). Only those ROIs with significant direct effects are shown here. The indirect model tested if likeness ratings might mediate the effects of the person-average or center-average on brain activity. Each point shows the proportion that each effect was mediated and the line shows the 95% confidence interval of the mediation. Lines above zero are significant and labeled with stars (*p* < 0.05, two-tailed).

### Face Familiarity

Next, we identified regions that were modulated by familiarity to each identity. We examined the effects of familiarity on brain activity using the average response times, more familiar faces should show faster response times on average (Supplementary Figure 5). Since the average likeness ratings for each identity were highly correlated with the response times (*r* = 0.57), the likeness ratings were excluded from this analysis. We found that the average RT for each identity significantly explained brain activity in the right V1 (t_5_=2.6, *p* < 0.05) and the left FFA (t_5_=2.8, *p* < 0.05). Seeing a more familiar face (i.e., face-identity with faster RT) was associated with lower brain activity in those two regions (right V1 and left FFA).

### Brain-Behavior Associations for Face Recognition and Person Identity

We regressed out-of-scanner likeness ratings and in-scanner response times onto brain activity to see if these measures were relevant for neural activity in our ROIs (Supplementary Figure 6). For response time, we found significant positive associations with brain activity (slower response times related to higher brain activity) in each of our ROIs (all *ps* < 0.05) except the right aFus. For likeness ratings, we found significant negative associations with brain activity (higher likeness ratings related to lower brain activity) in each of our ROIs (all *ps* < 0.05) except right/left V1 and left vATL.

**Supplementary Figure 5:**
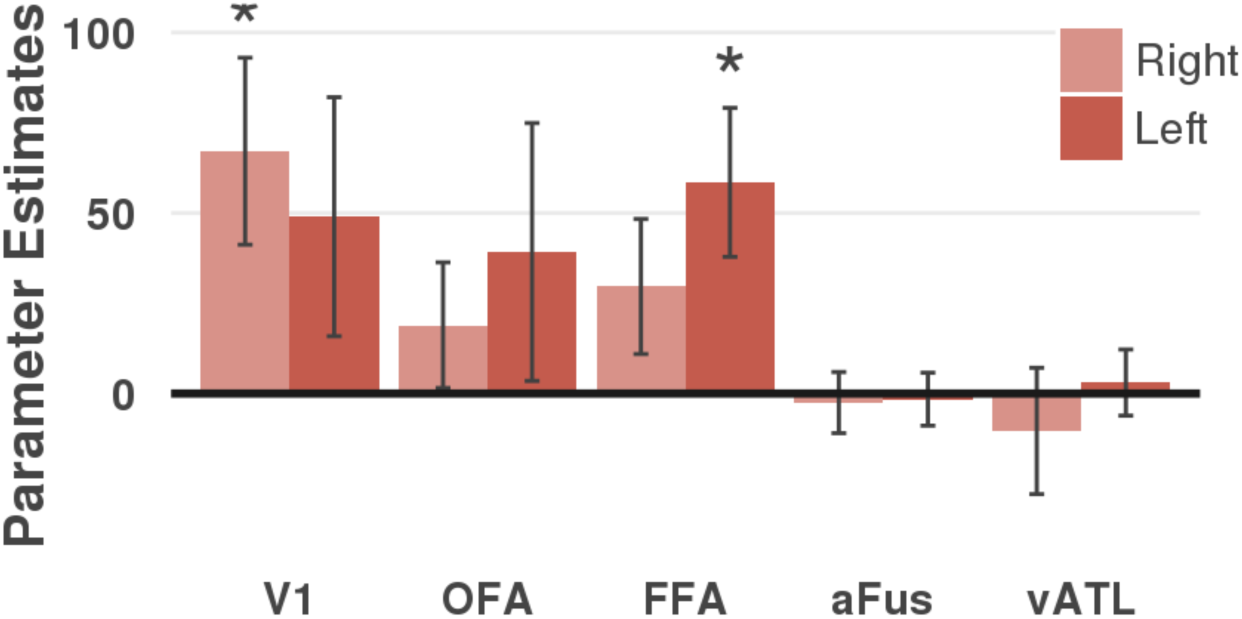
Bar plots showing brain activity (y-axis) for each ROI (x-axis) associated with the average response time for each identity. Brain activity is calculated as the average parameter estimates across participants and error bars show the associated standard error. Regions with asterisks are significant (p < 0.05).

**Supplementary Table 6a:** Listed are the peaks of significant ***brain activity positively associated with the likeness ratings*.** Peaks were found using AFNI’s 3dExtrema by setting a voxel-level threshold of Z > 1.96 (cluster corrected, *p* < 0.05) and a minimum distance of 12 mm between peaks. The last column provides the Z-Score for the Face > Scene contrast. Abbreviations: H, hemisphere; B, bilateral; R, right; L, left; BA = Brodmann Area.

| Brain region | H | BA | x | y | z | Z-Score | Face > Scene |
| --- | --- | --- | --- | --- | --- | --- | --- |
| **Likeness Ratings - Positive Associations** |  |  |  |  |  |  |  |
| *Cluster 1 (3,843 voxels)* |  |  |  |  |  |  |  |
| Postcentral sulcus | R | 3 | 48 | -24 | 42 | 4.21 | -0.55 |
| Posterior cingulate gyrus | R | 23 | 4 | -24 | 40 | 3.09 | -0.01 |
| Precentral gyrus | R | 6 | 56 | -6 | 46 | 3.39 | -1.57 |
| Precentral gyrus | R | 6 | 38 | -12 | 60 | 3.85 | 0.08 |
| Precentral sulcus | R | 6 | 18 | -20 | 64 | 4.11 | -1.95 |
| Precuneus gyrus | L | 5 | -8 | -50 | 66 | 4.17 | -1.73 |
| Superior frontal gyrus | L | 6 | -4 | -8 | 50 | 4.01 | -0.73 |
| Superior parietal gyrus | L | 7 | -38 | -48 | 68 | 4.36 | -1.21 |
| Superior parietal gyrus | R | 7 | 22 | -70 | 62 | 4.06 | 1.36 |
| *Cluster 2 (1,067 voxels)* |  |  |  |  |  |  |  |
| Lateral occipital gyrus | L | 18 | -10 | -98 | 24 | 3.67 | -1.00 |
| Precuneus gyrus | B | 23 | 0 | -66 | 20 | 3.69 | 1.18 |
| Superior parietal sulcus | R | 19 | 20 | -84 | 30 | 3.58 | -2.72 |
| *Cluster 3 (891 voxels)* |  |  |  |  |  |  |  |
| Inferior parietal gyrus | R | 40 | 58 | -54 | 40 | 3.75 | 0.74 |
| Superior temporal gyrus | R | 22 | 66 | -10 | -2 | 3.56 | -2.68 |
| Superior temporal gyrus | R | 22 | 68 | -34 | 14 | 4.66 | -0.30 |
| *Cluster 4 (819 voxels)* |  |  |  |  |  |  |  |
| Precentral sulcus | L | 6 | -32 | -16 | 60 | 3.06 | -1.33 |
| Supramarginal sulcus | L | 3 | -52 | -26 | 38 | 4.66 | 0.03 |
| *Cluster 5 (457 voxels)* |  |  |  |  |  |  |  |
| Cerebellum, V | B | NA | 0 | -60 | -12 | 3.87 | -0.08 |
| Cerebellum, VI | L | NA | -24 | -64 | -24 | 2.96 | 2.54 |
| Lingual sulcus | L | 19 | -28 | -58 | -2 | 4.27 | -2.76 |

**Supplementary Table 6b:** Listed are the peaks of significant ***brain activity negatively associated with the likeness ratings***. Peaks were found using AFNI’s 3dExtrema by setting a voxel-level threshold of Z > 1.96 (cluster corrected, *p* < 0.05) and a minimum distance of 12 mm between peaks. The last column provides the Z-Score for the Face > Scene contrast. Abbreviations: H, hemisphere; B, bilateral; R, right; L, left; BA = Brodmann Area.

| Brain region | H | BA | x | y | z | Z-Score | Face > Scene |
| --- | --- | --- | --- | --- | --- | --- | --- |
| **Likeness Ratings - Negative Associations** |  |  |  |  |  |  |  |
| *Cluster 1 (465 voxels)* |  |  |  |  |  |  |  |
| Fusiform gyrus (aFus) | L | 20 | -38 | -36 | -22 | -4.53 | 2.83 |
| Fusiform gyrus (aFus) | L |  | -36 | -20 | -20 | -3.35 | 1.09 |
| Inferior temporal gyrus | L | 20 | -48 | -48 | -20 | -3.28 | 2.40 |

**Supplementary Table 7:**
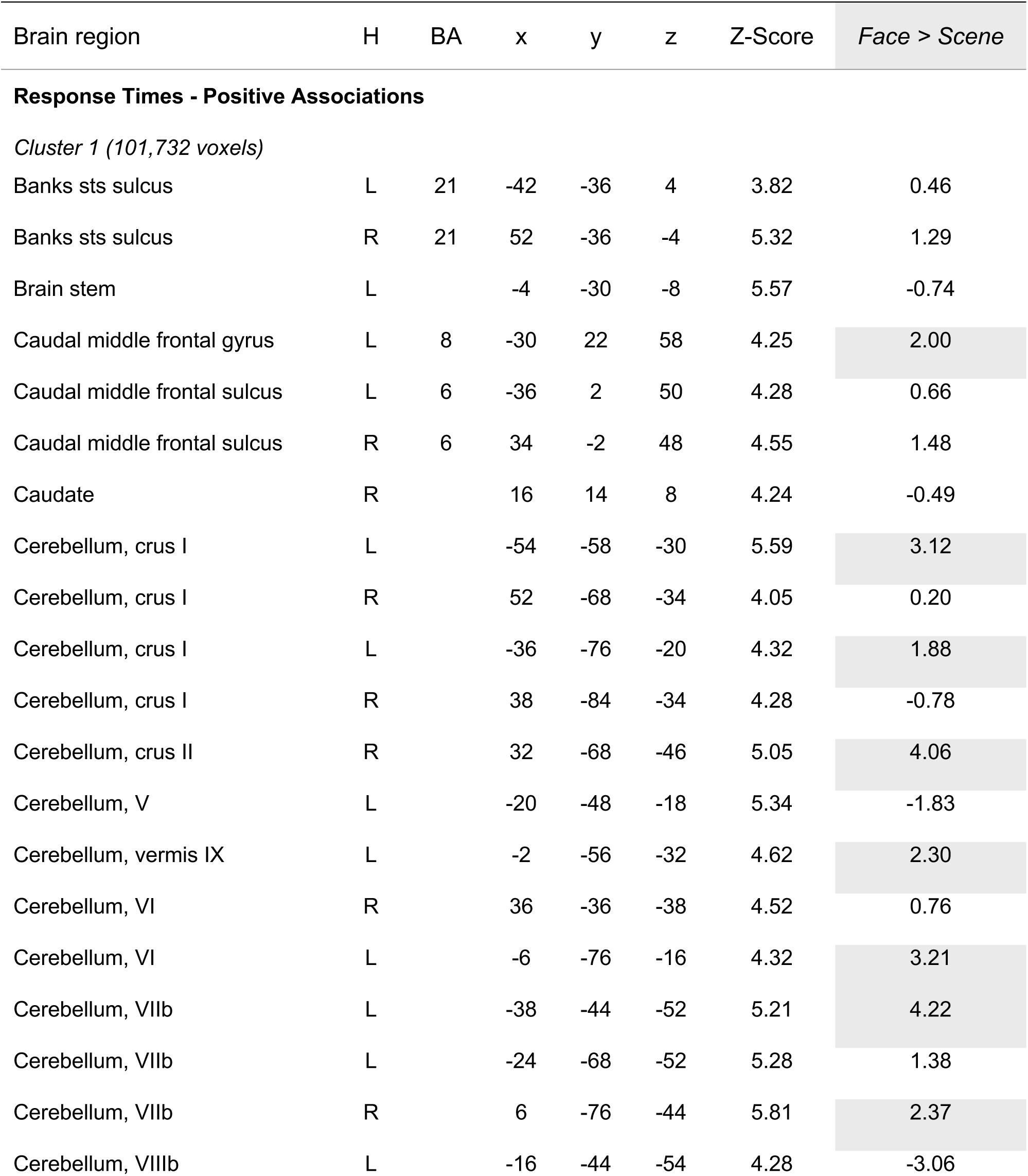
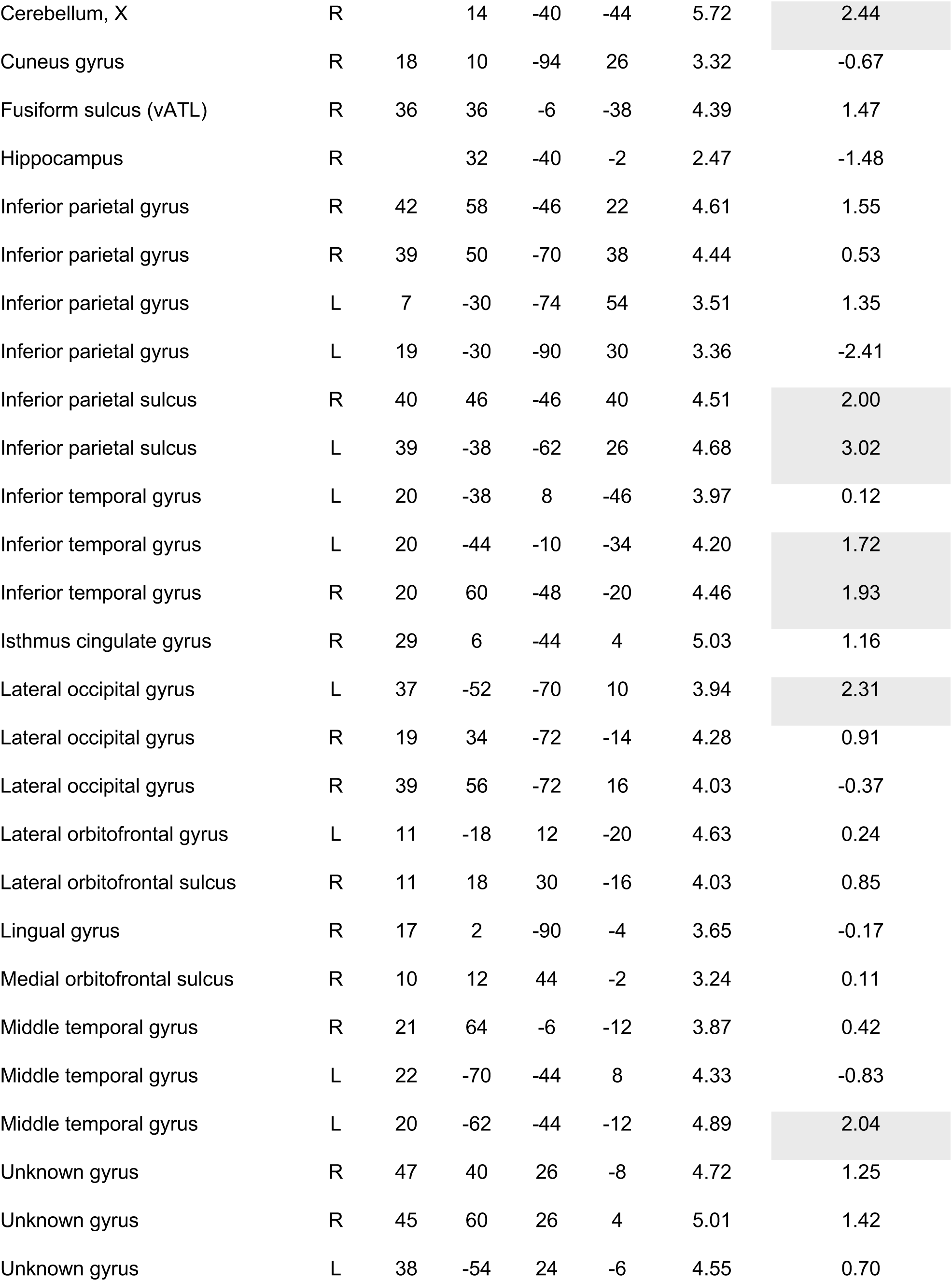
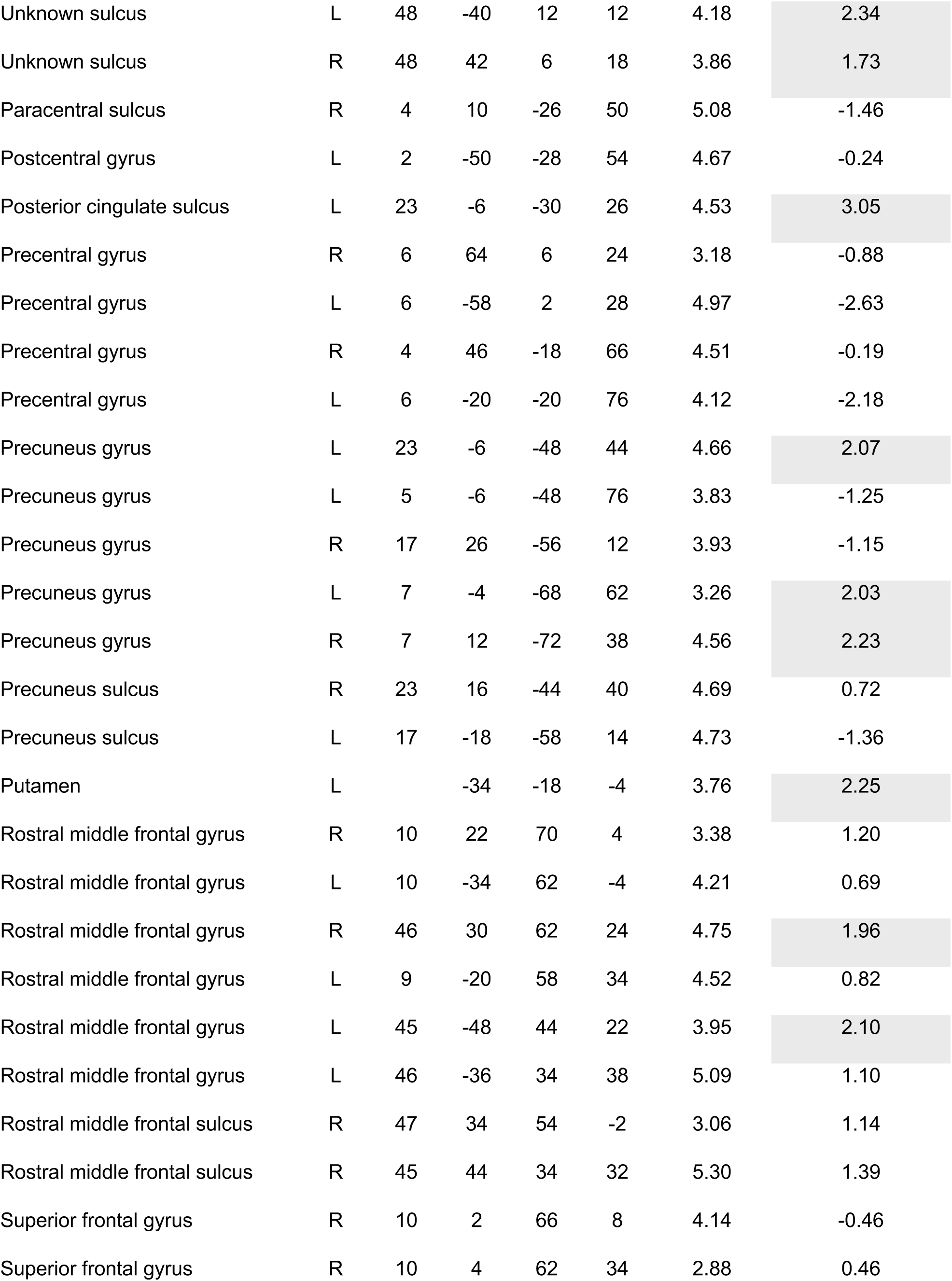
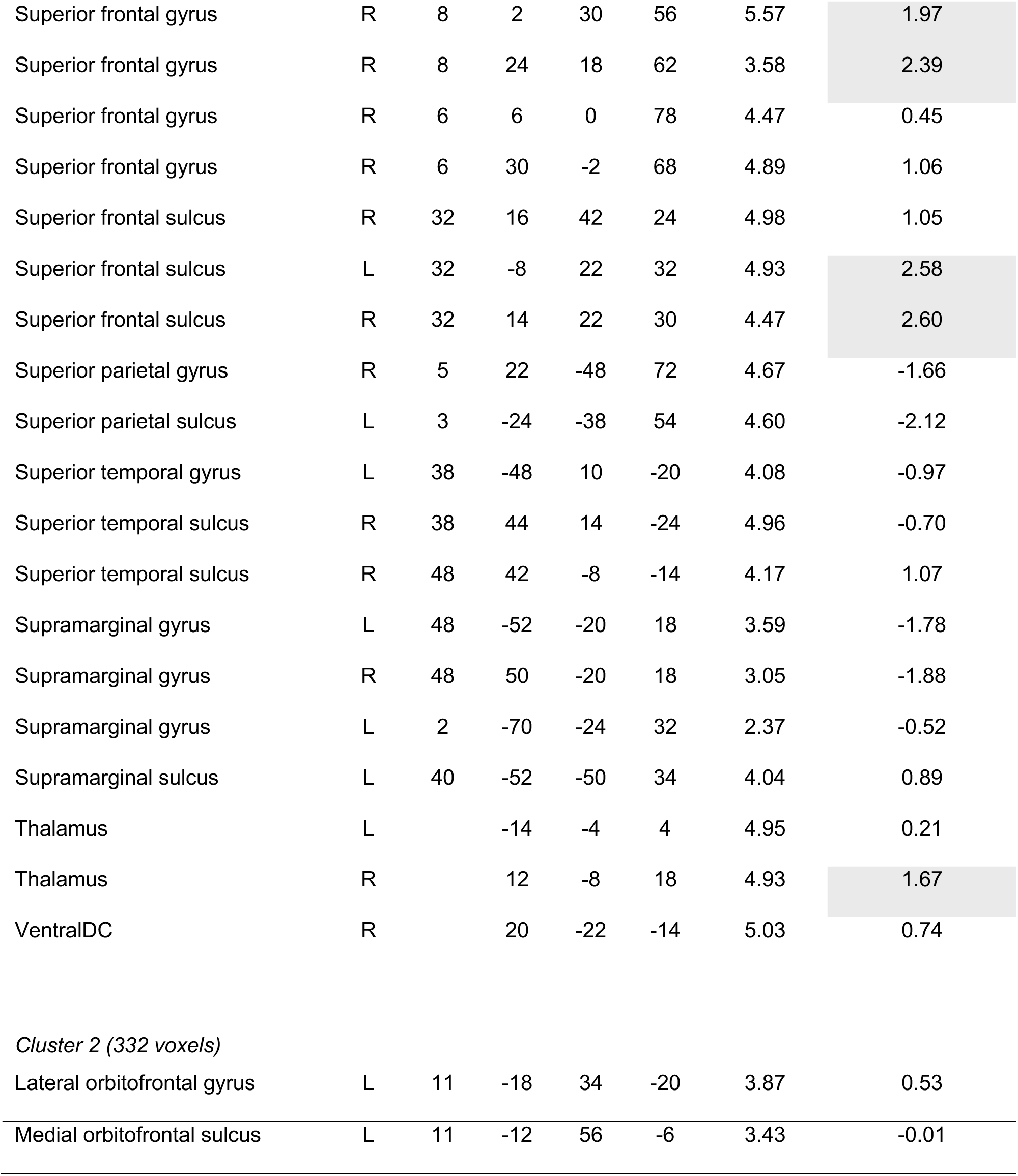
Listed are the peaks of significant ***brain activity associated with response time***. Peaks were found using AFNI’s 3dExtrema by setting a voxel-level threshold of Z > 1.96 (cluster corrected, *p* < 0.05) and a minimum distance of 12 mm between peaks. The last column provides the Z-Score for the Face > Scene contrast. Abbreviations: H, hemisphere; B, bilateral; R, right; L, left; BA = Brodmann Area.

| Brain region | H | BA | x | y | z | Z-Score | Face > Scene |
| --- | --- | --- | --- | --- | --- | --- | --- |
| **Response Times - Positive Associations** |  |  |  |  |  |  |  |
| *Cluster 1 (101,732 voxels)* |  |  |  |  |  |  |  |
| Banks sts sulcus | L | 21 | -42 | -36 | 4 | 3.82 | 0.46 |
| Banks sts sulcus | R | 21 | 52 | -36 | -4 | 5.32 | 1.29 |
| Brain stem | L |  | -4 | -30 | -8 | 5.57 | -0.74 |
| Caudal middle frontal gyrus | L | 8 | -30 | 22 | 58 | 4.25 | 2.00 |
| Caudal middle frontal sulcus | L | 6 | -36 | 2 | 50 | 4.28 | 0.66 |
| Caudal middle frontal sulcus | R | 6 | 34 | -2 | 48 | 4.55 | 1.48 |
| Caudate | R |  | 16 | 14 | 8 | 4.24 | -0.49 |
| Cerebellum, crus I | L |  | -54 | -58 | -30 | 5.59 | 3.12 |
| Cerebellum, crus I | R |  | 52 | -68 | -34 | 4.05 | 0.20 |
| Cerebellum, crus I | L |  | -36 | -76 | -20 | 4.32 | 1.88 |
| Cerebellum, crus I | R |  | 38 | -84 | -34 | 4.28 | -0.78 |
| Cerebellum, crus II | R |  | 32 | -68 | -46 | 5.05 | 4.06 |
| Cerebellum, V | L |  | -20 | -48 | -18 | 5.34 | -1.83 |
| Cerebellum, vermis IX | L |  | -2 | -56 | -32 | 4.62 | 2.30 |
| Cerebellum, VI | R |  | 36 | -36 | -38 | 4.52 | 0.76 |
| Cerebellum, VI | L |  | -6 | -76 | -16 | 4.32 | 3.21 |
| Cerebellum, VIIb | L |  | -38 | -44 | -52 | 5.21 | 4.22 |
| Cerebellum, VIIb | L |  | -24 | -68 | -52 | 5.28 | 1.38 |
| Cerebellum, VIIb | R |  | 6 | -76 | -44 | 5.81 | 2.37 |
| Cerebellum, VIIIb | L |  | -16 | -44 | -54 | 4.28 | -3.06 |
| Cerebellum, X | R |  | 14 | -40 | -44 | 5.72 | 2.44 |
| Cuneus gyrus | R | 18 | 10 | -94 | 26 | 3.32 | -0.67 |
| Fusiform sulcus (vATL) | R | 36 | 36 | -6 | -38 | 4.39 | 1.47 |
| Hippocampus | R |  | 32 | -40 | -2 | 2.47 | -1.48 |
| Inferior parietal gyrus | R | 42 | 58 | -46 | 22 | 4.61 | 1.55 |
| Inferior parietal gyrus | R | 39 | 50 | -70 | 38 | 4.44 | 0.53 |
| Inferior parietal gyrus | L | 7 | -30 | -74 | 54 | 3.51 | 1.35 |
| Inferior parietal gyrus | L | 19 | -30 | -90 | 30 | 3.36 | -2.41 |
| Inferior parietal sulcus | R | 40 | 46 | -46 | 40 | 4.51 | 2.00 |
| Inferior parietal sulcus | L | 39 | -38 | -62 | 26 | 4.68 | 3.02 |
| Inferior temporal gyrus | L | 20 | -38 | 8 | -46 | 3.97 | 0.12 |
| Inferior temporal gyrus | L | 20 | -44 | -10 | -34 | 4.20 | 1.72 |
| Inferior temporal gyrus | R | 20 | 60 | -48 | -20 | 4.46 | 1.93 |
| Isthmus cingulate gyrus | R | 29 | 6 | -44 | 4 | 5.03 | 1.16 |
| Lateral occipital gyrus | L | 37 | -52 | -70 | 10 | 3.94 | 2.31 |
| Lateral occipital gyrus | R | 19 | 34 | -72 | -14 | 4.28 | 0.91 |
| Lateral occipital gyrus | R | 39 | 56 | -72 | 16 | 4.03 | -0.37 |
| Lateral orbitofrontal gyrus | L | 11 | -18 | 12 | -20 | 4.63 | 0.24 |
| Lateral orbitofrontal sulcus | R | 11 | 18 | 30 | -16 | 4.03 | 0.85 |
| Lingual gyrus | R | 17 | 2 | -90 | -4 | 3.65 | -0.17 |
| Medial orbitofrontal sulcus | R | 10 | 12 | 44 | -2 | 3.24 | 0.11 |
| Middle temporal gyrus | R | 21 | 64 | -6 | -12 | 3.87 | 0.42 |
| Middle temporal gyrus | L | 22 | -70 | -44 | 8 | 4.33 | -0.83 |
| Middle temporal gyrus | L | 20 | -62 | -44 | -12 | 4.89 | 2.04 |
| Unknown gyrus | R | 47 | 40 | 26 | -8 | 4.72 | 1.25 |
| Unknown gyrus | R | 45 | 60 | 26 | 4 | 5.01 | 1.42 |
| Unknown gyrus | L | 38 | -54 | 24 | -6 | 4.55 | 0.70 |
| Unknown sulcus | L | 48 | -40 | 12 | 12 | 4.18 | 2.34 |
| Unknown sulcus | R | 48 | 42 | 6 | 18 | 3.86 | 1.73 |
| Paracentral sulcus | R | 4 | 10 | -26 | 50 | 5.08 | -1.46 |
| Postcentral gyrus | L | 2 | -50 | -28 | 54 | 4.67 | -0.24 |
| Posterior cingulate sulcus | L | 23 | -6 | -30 | 26 | 4.53 | 3.05 |
| Precentral gyrus | R | 6 | 64 | 6 | 24 | 3.18 | -0.88 |
| Precentral gyrus | L | 6 | -58 | 2 | 28 | 4.97 | -2.63 |
| Precentral gyrus | R | 4 | 46 | -18 | 66 | 4.51 | -0.19 |
| Precentral gyrus | L | 6 | -20 | -20 | 76 | 4.12 | -2.18 |
| Precuneus gyrus | L | 23 | -6 | -48 | 44 | 4.66 | 2.07 |
| Precuneus gyrus | L | 5 | -6 | -48 | 76 | 3.83 | -1.25 |
| Precuneus gyrus | R | 17 | 26 | -56 | 12 | 3.93 | -1.15 |
| Precuneus gyrus | L | 7 | -4 | -68 | 62 | 3.26 | 2.03 |
| Precuneus gyrus | R | 7 | 12 | -72 | 38 | 4.56 | 2.23 |
| Precuneus sulcus | R | 23 | 16 | -44 | 40 | 4.69 | 0.72 |
| Precuneus sulcus | L | 17 | -18 | -58 | 14 | 4.73 | -1.36 |
| Putamen | L |  | -34 | -18 | -4 | 3.76 | 2.25 |
| Rostral middle frontal gyrus | R | 10 | 22 | 70 | 4 | 3.38 | 1.20 |
| Rostral middle frontal gyrus | L | 10 | -34 | 62 | -4 | 4.21 | 0.69 |
| Rostral middle frontal gyrus | R | 46 | 30 | 62 | 24 | 4.75 | 1.96 |
| Rostral middle frontal gyrus | L | 9 | -20 | 58 | 34 | 4.52 | 0.82 |
| Rostral middle frontal gyrus | L | 45 | -48 | 44 | 22 | 3.95 | 2.10 |
| Rostral middle frontal gyrus | L | 46 | -36 | 34 | 38 | 5.09 | 1.10 |
| Rostral middle frontal sulcus | R | 47 | 34 | 54 | -2 | 3.06 | 1.14 |
| Rostral middle frontal sulcus | R | 45 | 44 | 34 | 32 | 5.30 | 1.39 |
| Superior frontal gyrus | R | 10 | 2 | 66 | 8 | 4.14 | -0.46 |
| Superior frontal gyrus | R | 10 | 4 | 62 | 34 | 2.88 | 0.46 |
| Superior frontal gyrus | R | 8 | 2 | 30 | 56 | 5.57 | 1.97 |
| Superior frontal gyrus | R | 8 | 24 | 18 | 62 | 3.58 | 2.39 |
| Superior frontal gyrus | R | 6 | 6 | 0 | 78 | 4.47 | 0.45 |
| Superior frontal gyrus | R | 6 | 30 | -2 | 68 | 4.89 | 1.06 |
| Superior frontal sulcus | R | 32 | 16 | 42 | 24 | 4.98 | 1.05 |
| Superior frontal sulcus | L | 32 | -8 | 22 | 32 | 4.93 | 2.58 |
| Superior frontal sulcus | R | 32 | 14 | 22 | 30 | 4.47 | 2.60 |
| Superior parietal gyrus | R | 5 | 22 | -48 | 72 | 4.67 | -1.66 |
| Superior parietal sulcus | L | 3 | -24 | -38 | 54 | 4.60 | -2.12 |
| Superior temporal gyrus | L | 38 | -48 | 10 | -20 | 4.08 | -0.97 |
| Superior temporal sulcus | R | 38 | 44 | 14 | -24 | 4.96 | -0.70 |
| Superior temporal sulcus | R | 48 | 42 | -8 | -14 | 4.17 | 1.07 |
| Supramarginal gyrus | L | 48 | -52 | -20 | 18 | 3.59 | -1.78 |
| Supramarginal gyrus | R | 48 | 50 | -20 | 18 | 3.05 | -1.88 |
| Supramarginal gyrus | L | 2 | -70 | -24 | 32 | 2.37 | -0.52 |
| Supramarginal sulcus | L | 40 | -52 | -50 | 34 | 4.04 | 0.89 |
| Thalamus | L |  | -14 | -4 | 4 | 4.95 | 0.21 |
| Thalamus | R |  | 12 | -8 | 18 | 4.93 | 1.67 |
| VentralDC | R |  | 20 | -22 | -14 | 5.03 | 0.74 |
| *Cluster 2 (332 voxels)* |  |  |  |  |  |  |  |
| Lateral orbitofrontal gyrus | L | 11 | -18 | 34 | -20 | 3.87 | 0.53 |
| Medial orbitofrontal sulcus | L | 11 | -12 | 56 | -6 | 3.43 | -0.01 |

### Prototype versus exemplar model

We assessed if an exemplar model provided a better fit to the brain data when compared to our previously applied prototype model. First, we examined how well the two models explained brain activity for faces overall, comparing the distances to the center average (prototype model) against the distances to the nearest faces in face-space (exemplar model) (Supplementary Figure 7a). None of the differences (prototype vs exemplar) in any region were significant, suggesting the two models explain the data equally well. Then, we examined how well the two models explained brain activity for faces of particular people, comparing the weighted distances to the person average (prototype model) against distance to the nearest faces of a person (exemplar model) (Supplementary Figure 7b). In this analysis also showed that none of the differences (prototype vs exemplar) in any region were significant, however the trend in the OFA, FFA, and aFus suggest that the prototype model on average provides a better fit than the exemplar model to brain activity.

**Supplementary Figure 6:**
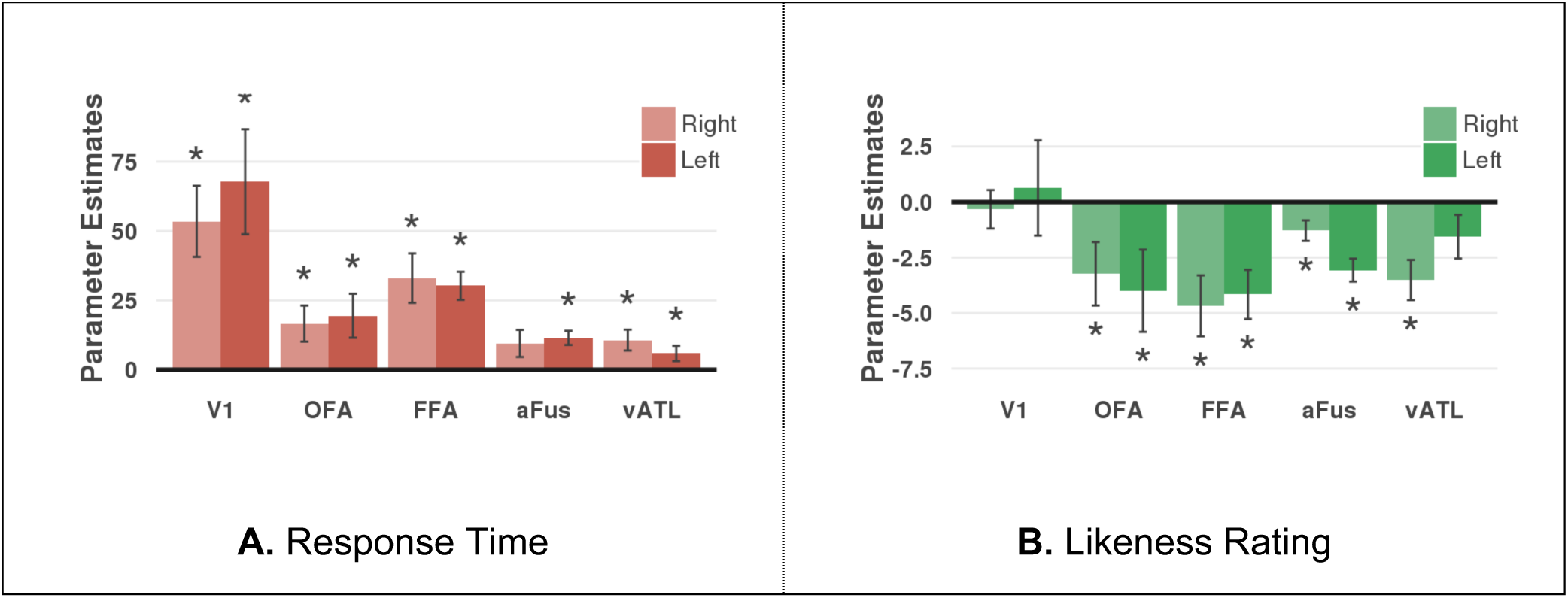
Bar plots show main effects of response time and likeness ratings on brain activity (t-statistics, y-axis) of the different ROIs (x-axis). The absolute t-statistics are used for better comparison between the two effects. All the response-time effects reflect positive t-statistics (slower response time leads to more brain activity) while all the likeness rating effects reflect negative t-statistics (the less a photo looks like that person results in more brain activity). Anything above the dotted red line indicates a significant effect (*p* < 0.05, two-tailed).

**Supplementary Figure 7:**
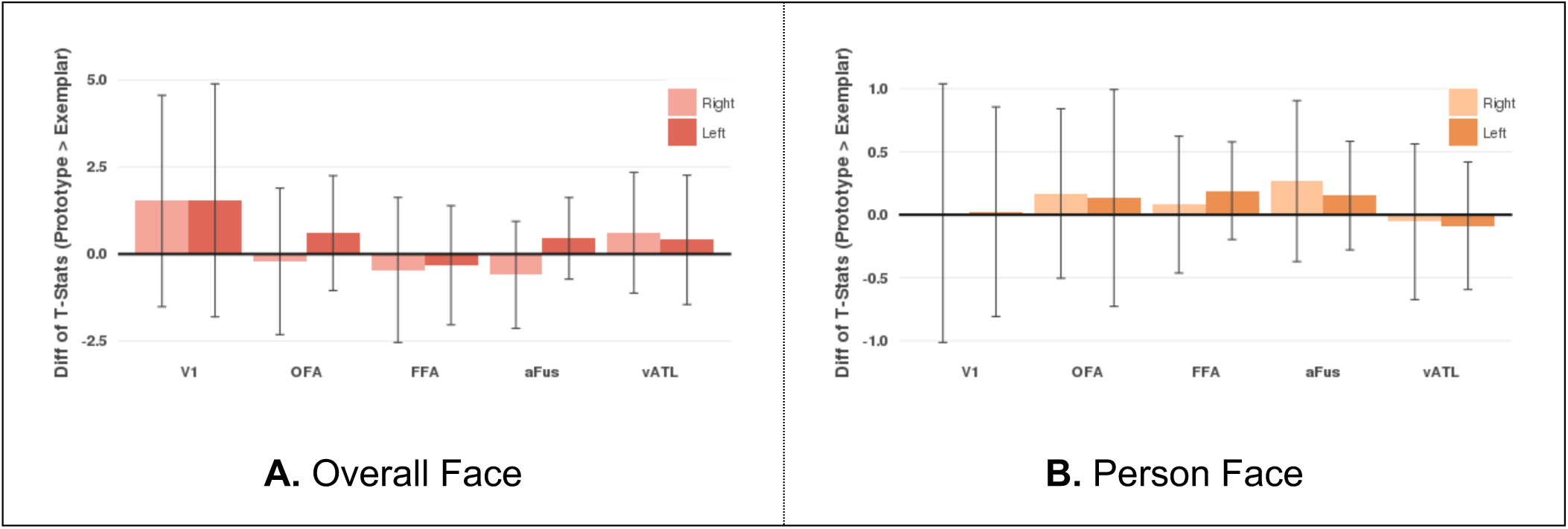
Differences in t-statistics of Prototype and Exemplar models (y-axis) are shown for each region (x-axis) when examining (a) faces overall and (b) faces for each person. Specifically, t-statistics represented fit of prototype (center and person averages) or exemplar (kNN to all and person faces) models to brain activity. Lighter color bars are for ROIs in the right hemisphere while darker color bars are for the left hemisphere. No bar was significantly different from zero.

